# Greenland’s thaw pushes the biodiversity crisis

**DOI:** 10.1101/2021.06.10.447623

**Authors:** Carolina Ureta, Santiago Ramírez-Barahona, Óscar Calderón-Bustamante, Pedro Cruz-Santiago, Carlos Gay-García, Didier Swingedouw, Dimitri DeFrance, Angela P. Cuervo-Robayo

## Abstract

Anthropogenic greenhouse gas emissions have led to sustained global warming over the last decades^1^. This is already reshaping the distribution of biodiversity across the world and can lead to the occurrence of large-scale singular events, such as the melting of polar ice sheets^2,3^. The potential impacts of such a melting event on species persistence across taxonomic groups – in terms of magnitude and geographic extent – remain unexplored. Here we assess impacts on biodiversity of global warming and melting of Greenland’s ice sheet on the distribution of 21,146 species of vascular plants and tetrapods across twelve megadiverse countries. We show that high global warming would lead to widespread reductions in species’ geographic ranges (median range loss, 35–78%), which are magnified (median range loss, 95–99%) with the added contribution of Greenland’s melting and its potentially large impact on oceanic circulation and regional climate changes. Our models project a decline in the geographical extent of species hotspots across countries (median reduction, 48–95%) and a substantial alteration of species composition in the near future (mean temporal dissimilarity, 0.26–0.89). These results imply that, in addition to global warming, the influence of Greenland’s melting can lead to the collapse of biodiversity across the globe, providing an added domino in its cascading effects.

---

Rising global temperatures are having negative impacts on biodiversity, increasing the risk of species extinctions across the world^4-8^. These negative impacts appear to be geographically widespread but are of serious concern for regions of high biodiversity fostering large numbers of restricted and endangered species^9^. These regions, known as biodiversity hotspots, are recognized as a critical priority for the conservation of the world’s biodiversity^10^. The most important hotspots are located within twelve megadiverse countries, which collectively harbor nearly two thirds (∼60%) of the Earth’s species of tetrapods and vascular plants^11^.

The current trends of greenhouse gas emissions and global warming are negatively impacting biodiversity within these hotspots, resulting in a substantial loss of species and the collapse of ecosystems within these regions^12-14^. Furthermore, if global warming continues unabated, there is the potential of catastrophic, large-scale singular events occurring, such as the disappearance of large areas of the Amazon rainforest or the melting of polar ice sheets. Accordingly, the Intergovernmental Panel on Climate Change has highlighted the need to incorporate such large-scale singular events into biodiversity risk assessments^14^. Even when the probability of occurrence of such events is not certain^12,13^, evaluating their possible consequences on biodiversity is a priority, yet these have been scarcely studied^9,15, 16^.

For instance, a substantial melting of Greenland’s ice sheets would generate an additional input of freshwater into the North Atlantic, leading to the weakening (or even complete shut-down) of the Atlantic Meridional Overturning Circulation (AMOC)^3^ – a key element of the global climate system. Recent evidence suggests that the rate of ice sheet loss over Greenland has accelerated over the last century^2,17-19^ and that the AMOC is currently the weakest it has been over the last millennium^18^. If these trends continue and a substantial melting of Greenland’s ice sheets occurs, this will induce further weakening of the AMOC – or even its collapse –, resulting in a significant rise in sea level^20,21^, and to alterations of regional climates^21-23^. These additional changes will build up the pressure on ecosystems and biodiversity across the world.

To date, a single study has evaluated the direct impacts of a weaker AMOC on biodiversity using an ecological niche modeling approach on amphibians^15^. As expected, the predicted impacts of global warming on amphibian decline are severe and widespread, and substantially enhanced by a weaker AMOC^12^. However, the extent and magnitude of these declines in species diversity across a broad range of taxonomic groups remain uncertain. Given that Greenland’s melting, and associated AMOC weakening, is expected to alter climatic zones around the world^22^, there is the need to assess the potential for this event to negatively impact biodiversity across a broader spectrum of terrestrial biodiversity, specifically by estimating alterations to species diversity and composition across the world’s biodiversity hotspots.

Here, we address this issue by constructing niche-based species distribution models (see **Materials and Methods**) for 21,146 species of tetrapods and vascular plants (amphibians, birds, mammals, reptiles, ferns, flowering plants, gymnosperms, and lycophytes) (**Extended Data figure 1**); the selected species are endemic to any of the twelve most megadiverse countries^11^ and are mostly located within biodiversity hotspots. By selecting this representative sample of species, we provide a general snapshot of the ecological impacts of unabated global warming and the melting of Greenland’s ice sheets. We quantify biodiversity loss and species’ vulnerability across countries and taxonomic groups through predicted changes in species diversity under five climate change scenarios: an unabated high-emission global warming scenario (RCP 8.5), and four variations of the RCP 8.5 generated by simulating different levels of freshwater release into the North Atlantic, starting in 2020^15,22^.

## Results and discussion

Our species distribution models for 21,146 tetrapods and vascular plants across twelve megadiverse countries project considerable changes in species richness (ΔSR) and composition (βSøR, see Methods) at three consecutive temporal horizons: 2030–2060, 2050–2080, and 2070– 2100 (hereafter referred to as 2030, 2050, and 2070, respectively). These time horizons should be interpreted as relative to the onset of the melting simulations in 2020^17,22^. As previously observed for amphibians^15^, the largest differences in species responses to climate change are observed between the global warming scenario (RCP 8.5) and any of the four melting scenarios. Given that differences between melting scenarios across the groups are small – even under the less drastic melting scenario we project drastic alterations to biodiversity –, we only show in the following the results obtained under scenarios RCP 8.5 and Melting 0.5 (**Supplementary table S1**), in order to gain clarity.

Our results agree with previous analyses showing a general decline in species richness under scenarios of global warming^9,24,25^. Here we show that melting has a large added impact on the observed patterns of biodiversity, probably as a result of induced major changes to temperature and precipitation^22^. The expected loss in species richness (negative ΔSR) appears to be more widespread with the added effects of melting than with global warming alone^12^, with some exceptions such as Indonesia, India, Philippines and China (**figure 1**). These results support the heterogeneous impacts of Greenland’s melting and associated AMOC weakening across the world, where melting can sometimes mitigate the impacts of changes induced by global warming^22^. However, excepting China, the above-mentioned countries have scarce biological data, thus geographical patterns of ΔSR should be interpreted with caution. Importantly, summarizing changes across plant and animal groups shows that there are differences in the geographic patterns of ΔSR between groups (**Supplementary figures S1–S2**). Overall, negative ΔSR are enhanced under melting scenarios but a contrasting pattern emerged for China, where the gain in species richness (positive ΔSR) increases by 2030 (**figure 1**); notwithstanding, even this trend of increasing species richness is reversed by 2050 and 2070.

**Figure 1.**
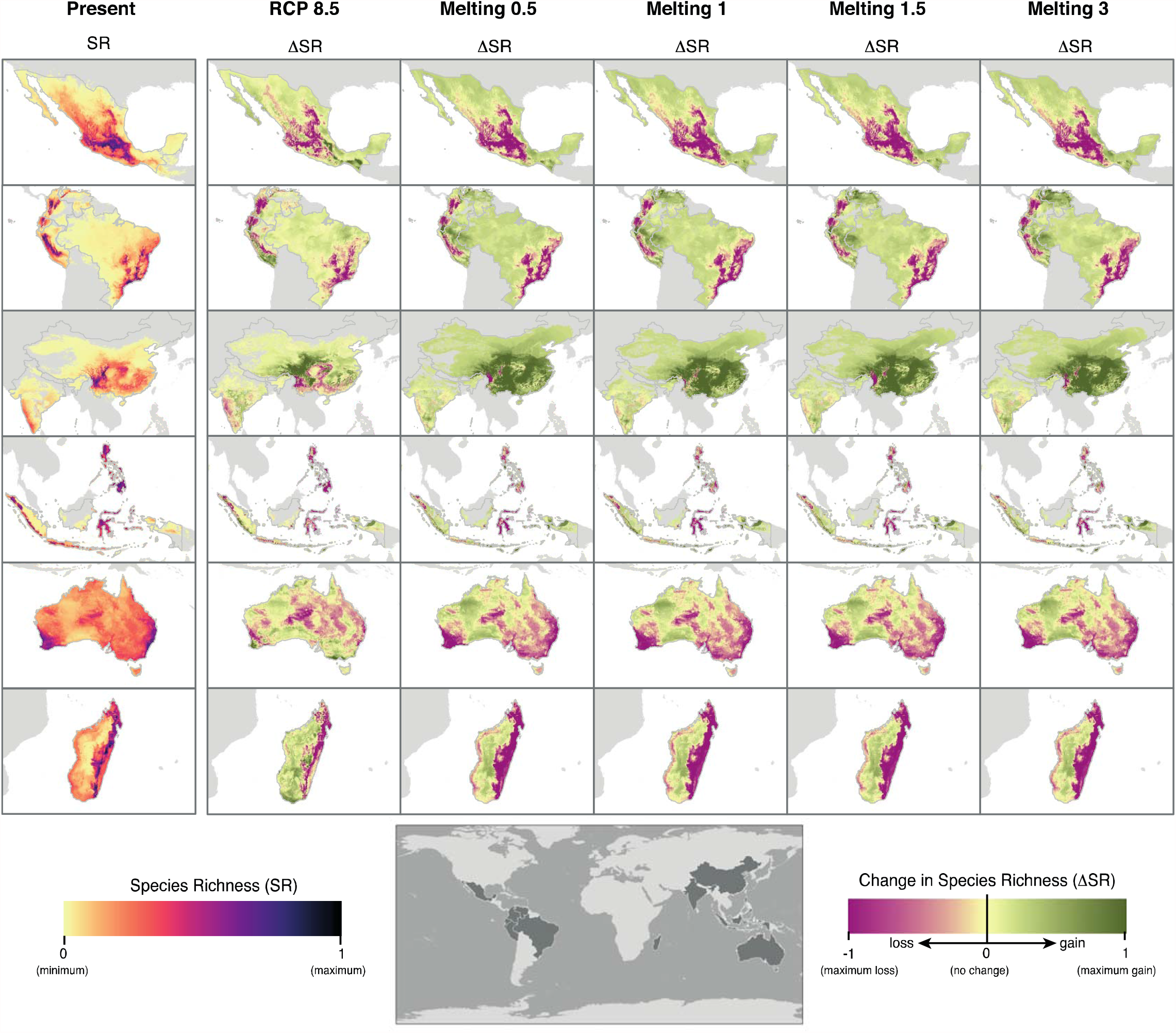
Geographic patterns of present-day species richness and temporal changes in species richness across twelve megadiverse countries. Estimates are based on species distribution models (SDMs) of vascular plants and tetrapods. Species richness (SR) was standardized to the range 0–1 within each country. Changes in species richness (ΔSR) are shown for 2030 under all scenarios. ΔSR was calculated as the difference in species richness between future scenarios and the present-day.

Based on our results, we suggest that on average, regions with more humid and temperate climates (*e*.*g*., mountain ecosystems) are predicted to be more vulnerable to climate change than regions with more seasonal and warmer climates^26-29^. These results coincide with the predicted reduction in Equatorial and warm temperate climates (A and C in the Köppen classification) across the world under melting scenarios^22^. For example, areas with positive ΔSR, both for tetrapods and vascular plants, are generally characterized by higher (or similar) mean annual temperatures and less (or similar) seasonality than areas with negative ΔSR, with the exception of Chinese tetrapods that show the opposite pattern (**Supplementary figure S3–S6)**. Differences in precipitation between areas of loss and gain of species richness are heterogeneous across countries. In general, areas with negative ΔSR for vascular plants appear to be more (or equally) humid (*i*.*e*., high annual precipitation and low seasonality) than areas with positive ΔSR; this pattern, however, appears to be less consistent, or even opposite, for tetrapods (**Supplementary figures S4, S6)**.

The vast majority of the species modeled are distributed within or around biodiversity hotspots^9^, including important mountain regions around the world. In this context, we show that the geographic extent of potential species hotspots (PSHs) across countries decreases – relative to their present extent – under global warming (median reduction, 48–88%) and are magnified with the added contribution of Greenland’s melting (median reduction, 74–95%) (**figure 2**; **Supplementary table S1**). Although most countries show decreasing trends of PSH extent, China shows marked increases under global warming (153–250%). This trend is reverted under melting scenarios, in which all twelve countries show sharp declines in the extent of PSH shortly after the onset of freshwater release into the North Atlantic (**Supplementary table S1**). In addition, our results indicate that PSHs would also be subjected to moderate to high changes in species composition (**Extended Data figure 2**), with a median temporal dissimilarity (β_SøR_) of 0.26–0.89 across scenarios. We observe an increased impact of Greenland’s melting – and the ensuing weaker AMOC – on species composition (**Supplementary table S2**); for instance, the median temporal dissimilarity is considerably lower under global warming alone (median dissimilarity, 0.18–0.51) than with the added contribution of melting scenarios (median dissimilarity, 0.20–0.90).

**Figure 2.**
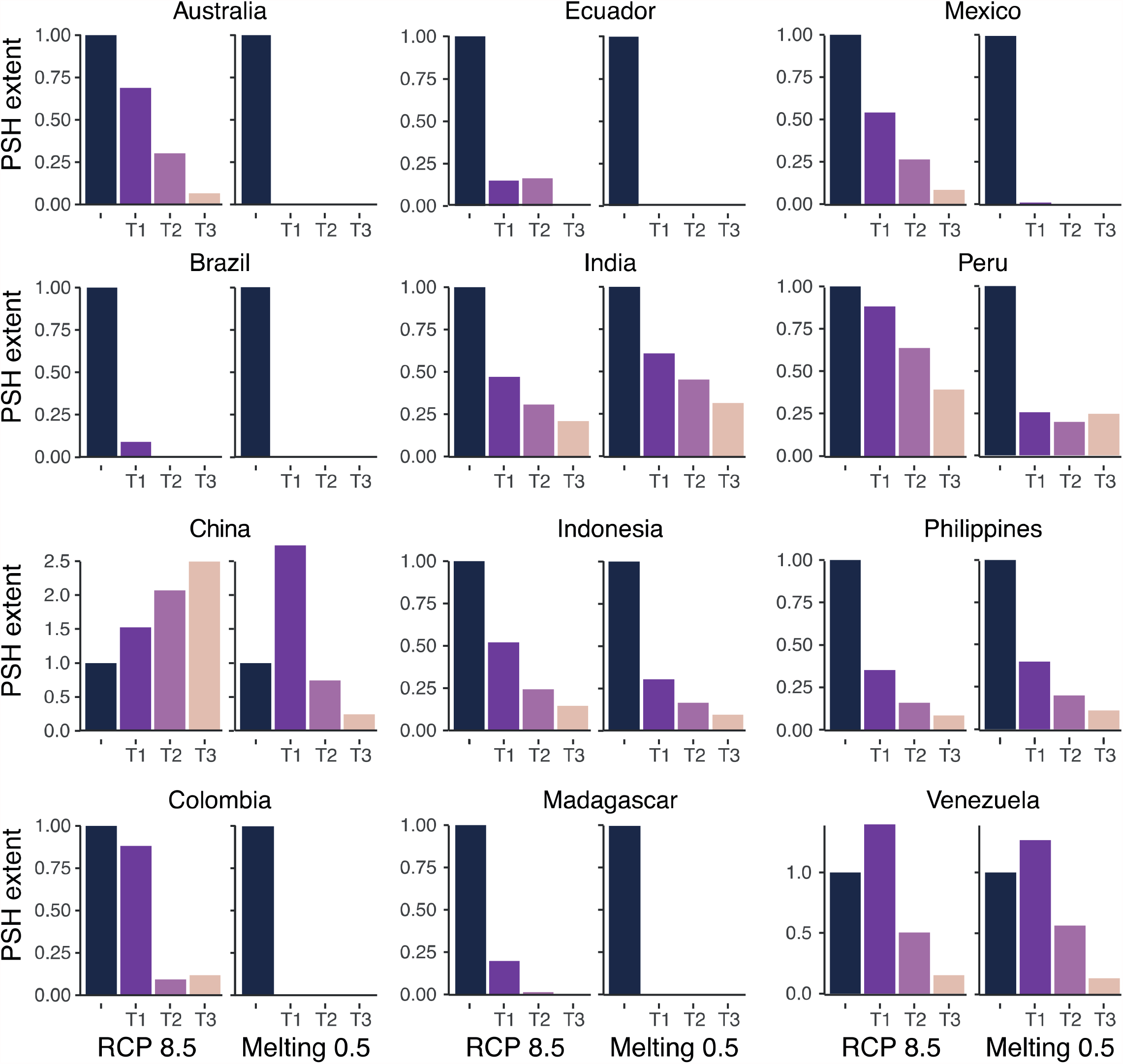
Temporal changes in the spatial extent of potential species hotspots (PSH) across twelve megadiverse countries. The extent of PSH was measured for each country separately as the number of pixels with a species richness (SR) higher than 0.6 × maximum SR. For each country, the extent of PSH was standardized relative to the present-day extent, where values greater than one indicate an expansion of PSHs and values of zero indicate the complete disappearance of PSHs. T1: 2030; T2: 2050; T3: 2070.

As expected, the potential species hotspots we defined coincide with globally important biodiversity hotspots^10^, which harbor climatically vulnerable species not found anywhere else in the world^30^. Thus, although our modeled species represent a small fraction of global diversity, the alterations to geographic extent of PSHs and the alteration of species composition is an alarming possibility. Based on our models, we suggest a dramatic decline and alteration of biodiversity across hotspots within a relatively short period of time (10–40 years; see figure S2 in Velasco et al.^15^) after the onset of Greenland’s melting, and the ensuing weakening of the AMOC. More than twenty years ago, Myers et al.^10^ estimated that effectively protecting these biodiversity hotspots, collectively encompassing less than 2% of the Earth’s surface, would translate into the protection of 44% of vascular plants and 35% tetrapods. However, our results indicate that, even if climate mitigation and ecosystem protection and restoration take place, the world’s biodiversity hotspots are highly vulnerable in the face of tipping points pushing the climate system into a new state^16,31^.

The projected reduction in species richness and alteration to species composition highlight the threat for biodiversity posed by global warming and the additional contribution of Greenland’s melting^15^. Furthermore, the declines in species richness and alteration to species composition are associated with projected reductions of the geographic ranges of individual species, which are magnified under melting scenarios (**figure 3**). We project that under global warming alone, half of the vascular plant and tetrapod species will experience range reductions of at least 31–83%, relative to present-day distributions (**Extended Data figure 3; Supplementary table S3–S4**), with reductions across individual countries ranging from 10–38% (Peru) to 54– 79% (Brazil). However, we estimate range expansions in more than half of the species in China, Colombia, and Venezuela (median expansion, 102–120%), albeit only for 2030 (**Supplementary table S4**). Nearly all range expansions are reversed towards range reductions with the added contribution of Greenland’s melting (median range loss, 95–99%). Under melting scenarios, species with a projected extreme to complete loss of geographic ranges (more than 80% reduction) increase in number relative to the RCP 8.5 scenario (**Extended Data figure 4**). More specifically, species ranges for the eight taxonomic groups are reduced drastically after the onset of freshwater release (**Extended Data figures 3–4)**, with a median range reduction of 58–99% by 2030, increasing to 67–100% by 2070 (**Supplementary tables S3**); these projections mean that in the worst-case scenario, half of the species will suffer complete range reductions.

**Figure 3.**
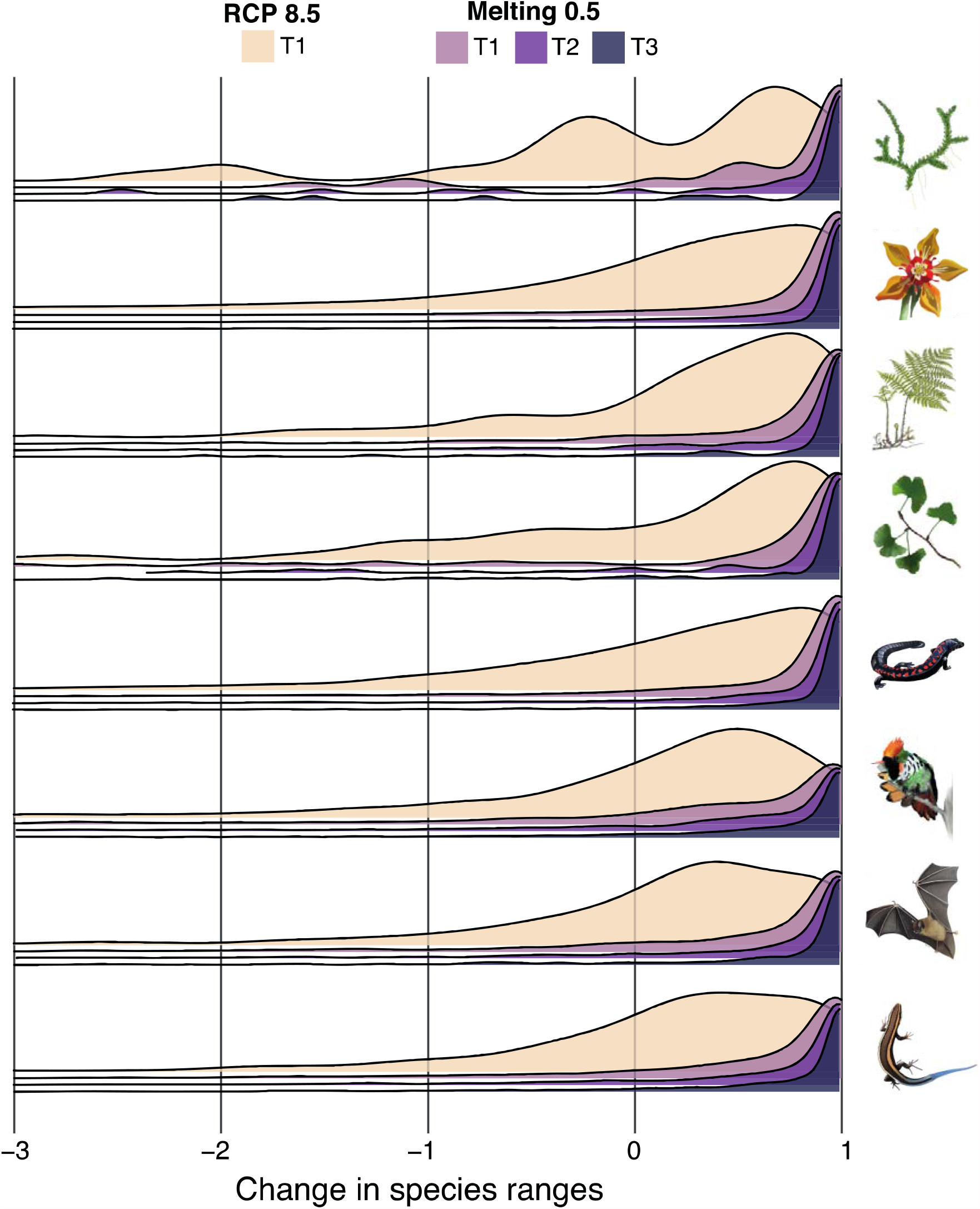
Changes in the size of the distribution range of species of vascular plants and tetrapods. Estimates of range size were based on species distribution models (SDMs). Range sizes were standardized relative to the present-day species range size: positive values indicate range reductions (1 means complete loss) and negative values indicate range expansions. Estimates of range size were aggregated by taxonomic group and visualized using density plots for T1: 2030 under the RCP 8.5 scenario and for T1: 2030, T2: 2050, and T3: 2070 under the Melting 0.5 scenario. Groups of species are indicated by the inset illustration (from top to bottom): lycophytes, flowering plants, ferns, gymnosperms, amphibians, birds, mammals, and reptiles.

It is safe to assume a direct relationship between the loss of suitable areas and species’ extinction risks^32^, where species with reduced geographic ranges are more prone to extinction than wide-ranging species. In the present case, the complete disappearance of suitable areas for species within megadiverse countries, which is exacerbated by melting, is of paramount concern because these losses entail high probabilities of species being fully extirpated from their native range and going extinct globally. Our results indicate that, in general, plant species have slightly higher risks of extinction (median range loss, 37–100%) than animal species (median range loss, 30–96%) across all scenarios (**Supplementary table S5**). Our models show a complete disappearance of climatically suitable areas for 1,239–4,483 species (6–21% of the total) under global warming, which increases to 7,728–10,312 species (36–49% of the total) with the added contribution of Greenland’s melting (**Extended Data figure 4, Supplementary figures S7–S8**).

The general pattern of complete range reduction is ubiquitous across countries, but with some variation (**figure 4, Supplementary table S6**). For instance, Brazil and Australia show the highest proportion of species (relative to the species modeled per country) projected with complete range loss (proportion of species, 8–60% and 7–59%, respectively), which is consistent with substantial alterations to regional climates predicted under melting scenarios^22^. On the other hand, India and Philippines show the lowest proportion of species with complete range loss (proportion of species, 3–18% and 2–19%, respectively) (**Supplementary table S6**). The estimated changes in our evaluated variables of mean annual temperature and annual precipitation under the different climate models cannot account for the spatial and temporal heterogeneity observed in species responses across countries (**Supplementary figures S9–S11**); this despite that these variables (mean annual temperature and annual precipitation) are two of the most important contributors to our species distribution models (**Supplementary figure S12– S13**). This highlights the idiosyncratic response of species to climate change, yet our results suggest that high global warming and ice sheet melt can have an overarching impact on biodiversity and the climate system^15,22^, leading to worldwide drastic alterations to climate and biodiversity loss.

**Figure 4.**
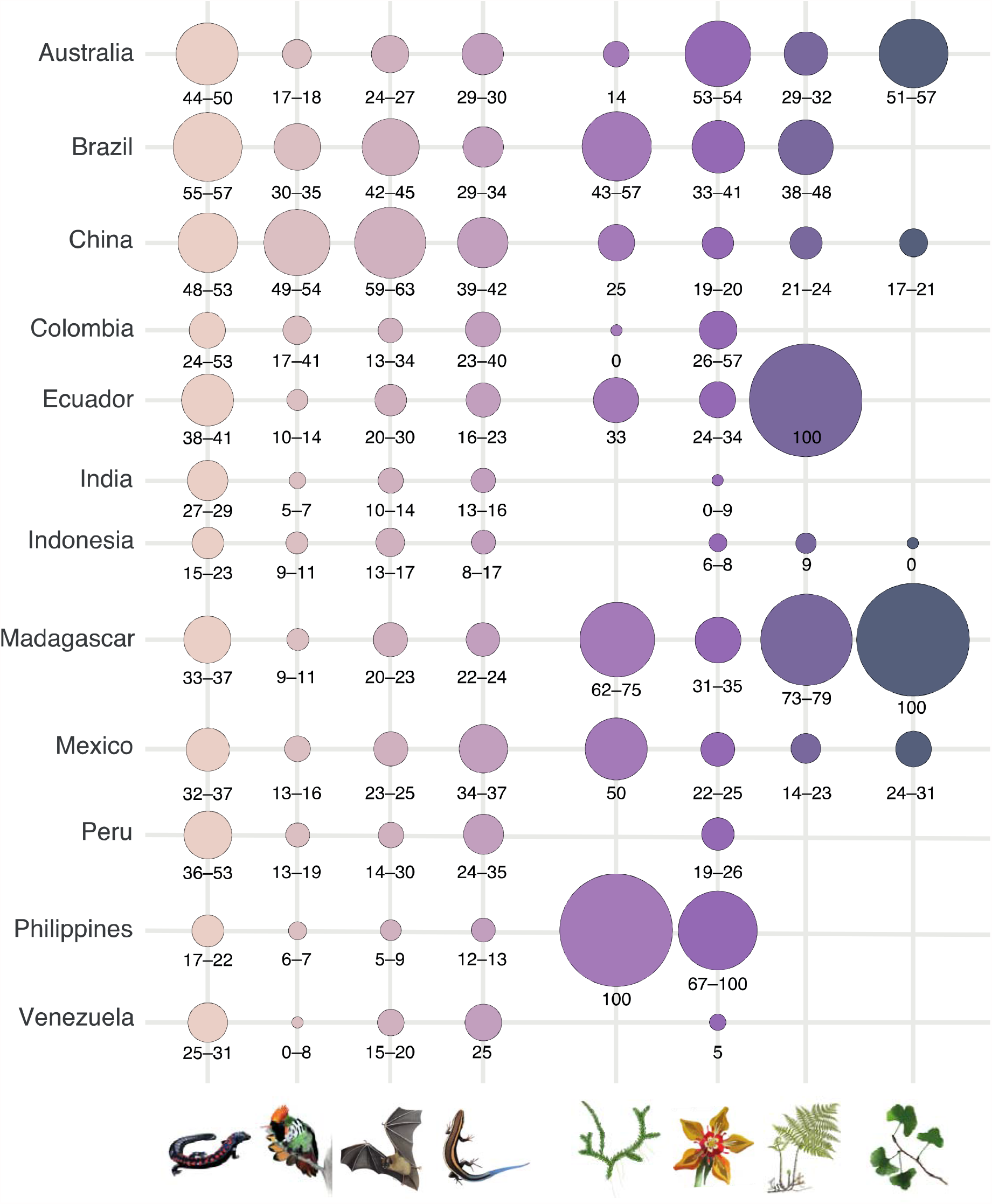
Proportion of species of vascular plants and tetrapods with complete range loss across twelve megadiverse countries. The proportion of species (%) with complete range loss was estimated relative to the total number of species modelled for each group and country. Numbers below circles indicate the range of values estimated for 2030 across the four Melting scenarios. Range size estimation was based on species distribution models and were standardized relative to the present-day species range size. Groups of species are indicated by the inset illustration (from left to right): amphibians, birds, mammals, reptiles, lycophytes, flowering plants, ferns, and gymnosperms.

The projected range losses would have a considerable impact for the future of the worlds’ biodiversity and, in the case of South American countries, further increase the global concerns on the region’s deforestation trends^14,33^, an alternative large-scale singular event. On the other hand, island countries, such as Indonesia and the Philippines, are projected to have smaller reductions in species ranges. In part, we think this is the result of a lower number of modeled species, which we believe to be related to a lack of publicly available biodiversity data in these countries, rather than an inherent lower climatic vulnerability^15,22^. Data limitations in some of these countries limit the assessment capabilities of species vulnerability, thus hindering conservation and mitigation planning^34^. Sources of uncertainty in prediction, including modelling uncertainty, need to be taken into account when assessing vulnerability of regional biodiversity under climate change^35-37^ and to define conservation priorities^38^. In this respect, although the species distribution modelling is based on a single global circulation model and inter-model variability is not considered, the resulting species models take into account modelling uncertainty by relying on an ensemble approach with seven different algorithms. Thus, our species distribution models allowed us to assess for the first time the probable consequences of Greenland’s melting on biodiversity.

Based on our results, we suggest an additional domino effect might be triggered in the potential cascading effects of anthropogenic climate change, where biodiversity will be crossing a tipping point in response to Greenland ice sheet thawing. In this context, alterations to species composition and decreasing species diversity are expected to have further cascading effects on biological interactions and ecosystem functioning, further increasing the probability of species’ extinction^37^. In this context, we show that on average, vascular plants are expected to be the most vulnerable (**Extended Data figure 3**), but this might partially be the result of the small number of species modelled for ferns, gymnosperms, and lycophytes. Nonetheless, flowering plants are by far the most abundant group in our dataset – 15,162 species – and apart from the three aforementioned groups, it is the most vulnerable group in terms of complete loss of species geographic ranges. The fact that flowering plants are projected to suffer substantial negative impacts increases the concerns about the vulnerability of biodiversity in general, due to possible future alterations to ecological interactions and ecosystem functioning^8, 39,40^. In turn, the two most vulnerable tetrapod groups are amphibians and reptiles^15,41-43^, yet mammals and birds also appear to be highly vulnerable in some countries (**figures 4**). These groups of animals may be further impacted due to their dependence on unaltered and diverse forest ecosystems. In this context, the climatic vulnerability of flowering plants – which are the ecological basis of most terrestrial ecosystems – can potentially increase further due to the collapse of tetrapod diversity, which includes many pollination and dispersal vectors. The collapse of flowering plant diversity will likely increase the extinction risks of other ecologically linked groups^8,39^, even when these appear to be less vulnerable to climate change, such as birds in Andean countries (**figure 4**).

Based on the projected changes to the geographic range of thousands of animal and plant species, we suggest a likely tipping point in the collapse of biodiversity across the world’s most megadiverse countries in response to unrelenting global warming. These impacts would be magnified and be reached relatively quickly (10–40 years), if a substantial amount of freshwater from Greenland’s ice sheets is released into the North Atlantic. In light of recent observations of a substantial ice sheet loss and of the AMOC being currently at its weakest point in millennia^17,18^, our projections provide reasons of major concern for the future of species across major biodiversity hotspots. The effects of global warming and Greenland’s melting would be reflected in substantial reductions to species diversity and alteration to species composition. If current trends of climate change continue – resulting from unrelenting greenhouse gas emissions – these effects have the potential to lead to substantial alterations to ecological interactions and ecosystem functioning.

## Materials and Methods

### Species occurrence records

We compiled data on the distribution of species of four tetrapod and four vascular plant groups (amphibians, reptiles, birds, mammals, lycophytes, ferns, gymnosperms, and flowering plants; **Supplementary Methods**). Species occurrence records were obtained from the Global Biodiversity Information Facility (GBIF)^44^, the International Union of Conservation of Nature (IUCN)^45^, and BirdLife^46,47^. For vascular plants, we used geographic occurrence data obtained from the Global Biodiversity Information Facility by querying all records under ‘Tracheophyta’ (we only considered ‘Preserved Specimens’ in our search). We taxonomically homogenized and cleaned the occurrence records following the procedures described in Ramírez-Barahona et al.^48^ using Kew’s Plants of the World database^49^ as the source of taxonomic information. For countries in which data for vascular plants were scarce or absent (*e*.*g*., India), we complemented occurrence information with IUCN polygons from the IUCN (although IUCN data for plants remains limited).

We explored the possibility of using occurrence records from GBIF for tetrapods, but data for megadiverse countries were scarce. Consequently, we decided to use the distribution polygons provided by the IUCN for amphibians, reptiles and mammals (terrestrial and freshwater species)^45^, and the distribution polygons provided by BirdLife^46^. This decision was based on the fact that ecological niche modeling using IUCN polygons has been proven to give robust results^15^. For the IUCN polygons, we retained species that have been categorized as “extant”, “possibly extinct”, “probably extant”, “possibly extant”, and “presence uncertain”, discarding species considered to be “extinct”. In addition, we did not model species reported by the IUCN as “introduced”, “vagrant”, or those in the “assisted colonization” category; for mammals and birds we only considered the distribution of “resident” species.

Depending on the taxonomic group, and given the information available, we used different approaches to identify species endemic to any of twelve megadiverse countries: Australia, Brazil, China, Colombia, Ecuador, India, Indonesia, Madagascar, Mexico, Peru, Philippines, and Venezuela. For birds, we used BirdLife to identify species listed as “breeding endemic” and then choose the corresponding IUCN polygons. To identify endemic species, we used a 0.08333° buffer around each country to select the IUCN polygons that fall completely within the country limits. We converted all selected species polygons into unique records at a 5 min resolution (approximate 10 km at the equator). For the point occurrence data for vascular plants, we identified endemic species as those with all occurrence records restricted to any given megadiverse country. We only modeled species with at least 25 unique records at a 5 arc-minute resolution (approximate 10 km at the equator). In many cases, the processing of the IUCN polygons for tetrapods resulted in species with thousands of occurrence records. In these cases, we randomly chose a maximum of 500 hundred records per species.

### Climatic data

We used the bioclimatic variables available at WorldClim v.2^49^ as the baseline (present-day) climatic conditions (1970–2000). For the future scenarios, we used bioclimatic variables derived the IPSL-CM5-LR ocean-atmospheric model under five scenarios: i) the high-emissions RCP 8.5 W/m^2^ scenario (2006–2100); and ii) Melting scenarios consisting of four different experiments of freshwater discharge into the North Atlantic from Greenland’s meltwater (see DeFrance et al.^23^ for a details). The four melting scenarios are equivalent to a sea level rise of 0.5, 1, 1.5, and 3 meters above current levels; these are named accordingly: Melting 0.5, Melting 1, Melting 1.5., and Melting 3. We obtained debiased bioclimatic variables^23^ under the five future scenarios for three consecutive time horizons: 2030–2060; 2050–2080; and 2070– 2100.

### Ecological niche modeling

At their most basic, the algorithms used to construct species distribution models relate species occurrence records with climatic variables in order to create a climatic profile that can be projected onto other time periods and geographic regions^50^. The resulting models have proven useful in evaluating the impacts of climate change on biodiversity, and to identify varying levels of vulnerability among species^35,36,38^. Here, we employed a multi-algorithm (ensemble) approach to construct species distribution models as implemented in the “biomod2” package^51^ in R^52^. The underlying philosophy of ensemble modeling is that each individual model carries a true “signal” about the relationships the model is aiming to capture, and some “noise” created by errors and uncertainties in the data and the model structure^35,36^. By combining models created with different algorithms, ensemble models aim at capturing the true ‘signal’ while controlling for algorithm-derived model differences; therefore, model uncertainty is accounted for during model construction.

Prior to modelling, we reduced the number of bioclimatic variables per species by estimating collinearity among present-day bioclimatic variables. We employed the ‘corrSelect’ function of the package fuzzySim^53^ in R^52^, using a Pearson correlation threshold of 0.8 and variance inflation factors as criteria to select variables. We used seven algorithms with a good predictive performance (evaluated with the TSS and ROC test): Maxent (MAXENT.Phillips), Generalized Additive Models (GAM), Classification Trees Analysis (CTA), Artificial Neural Networks (ANN), Surface Range Envelope (SRE), Flexible Discriminant Analysis (FDA), and Random Forest (RF). As occurrence datasets consisted of presence only data, for each model, we randomly generated 10,000 pseudo-absences within the model calibration area; we set the prevalence to 0.5 in order to give presences and absences the same importance during the calibration process. For each species we selected a calibration area (or M)^50^ using a spatial intersection between a 4º buffer around species occurrences and terrestrial ecoregions^54^ (**Supplementary figure S14**). By doing this, we incorporated information about dispersal and ecological limitations of each species into the modelling^50^.

We calibrated each algorithm using a random sample of 70% of occurrence records and evaluated the resulting models using the remaining 30% of records. To validate the models, we used the True Skill Statistics (TSS) and performed 10 replicates of every model, providing a ten-fold internal cross-validation. As a way to deal with uncertainty, we constructed the ensemble models (seven algorithms × ten replicates) using a total consensus rule, where SDMs from different algorithms are assembled using a weighted mean of replicates with an evaluation threshold of TSS > 0.7. We encountered problems during ensemble construction for some widespread bird and reptile species, where all seven algorithms had a performance below the specified evaluation threshold; consequently, for these species we reduced the evaluation threshold to TSS > 0.6. In addition, due to modelling issues in some insular species we changed the calibration area (M) to the entire country.

For each species, we used the resultingh ensemble models to project the potential distribution of each species under both current and future climatic conditions. The distribution of validation statistics (TSS, ROC) for the ensemble SDMs are shown in **Supplementary figure S15**. In sum, we obtained sixteen ensemble probability maps per species, one for the present day and fifteen for the five scenarios and the three different time horizons; we also summarized the mean coefficient of variation – reflecting model uncertainty – across species present in each grid-cell. Finally, we examined the frequency of different bioclimatic variables as the most important contributing variables during model construction; for every species, we retrieved the two variables with the largest model contribution.

### Species richness, temporal dissimilarity, and geographic ranges

We converted ensemble probability maps into binary maps of presence/absence using the TSS threshold. Species richness (SR) was estimated as the sum of species present in each grid-cell; we generated 16 species richness maps corresponding to the present-day and the four future scenarios at each of the three temporal horizons. We used these maps to estimate species richness (SR) across space and to calculate the change in species richness (ΔSR) through time. We assumed full dispersal ability of species in all analyses, meaning that all suitable areas in the future are assumed to have the same probability of being occupied by the species irrespective of the distance to the present-day distribution. We standardized species richness per country and estimated the change in species richness (ΔSR) across grid-cells by subtracting the estimated SR in the future from the current SR; negative ΔSR values indicate species’ loss and positive ΔSR indicate species’ gain.

We defined Potential Species Hotspots (PSH) within each country as those grid-cells with the highest levels of species richness. For this, we used a threshold of 60% of the highest level of species richness observed in the present-day within each country. Considering only those grid-cells with a richness value above the threshold, we estimated the geographic extent of PSHs across time periods and scenarios and estimated reductions and expansions of PSHs relative to their present-day extent; we tested additional thresholds to define and quantify the extent of PSHs (**Supplementary Methods**).

We estimated the change in species composition through time using the Sørensen pairwise dissimilarity index (β_SøR_), which estimates the dissimilarity in species composition between two sites and incorporates both turnover and differences in species richness among sites. For this, we estimated dissimilarity between the present-day and each of the three temporal horizons at each spatial location within PSHs; we computed temporal dissimilarity across all PSHs and scenarios (**Supplementary Methods**). In the context, the observed temporal dissimilarity reflects two main patterns of varying composition under climate change scenarios: (i) the replacement of present-day species by ‘new’ species within sites; and (ii) the loss (or gain) of species resulting in nested species assemblages. Values of temporal β_SøR_ approaching one are indicative of higher dissimilarity between the present-day species composition and the future projected composition within sites, and values approaching zero are indicative of few temporal changes in composition.

Finally, in order to approximate the vulnerability of individual species to climate change, we estimated the temporal changes in the extent of the potential distribution area (range size) for every species relative to the present-day distribution. This allowed us to estimate the degree to which the projected geographic range for each species would be reduced (or expanded) under different scenarios. In turn, we estimated the proportion of species (by country and group) projected to have a complete loss of potential distribution area in the future. Finally, we compared the change in species range sizes between scenarios by defining five categories: (i) gain, species with positive changes in range size; (ii) moderate loss, species with changes in range size between 0 and -0.45; (iii) severe loss, species with changes in range size between -0.45 and -0.8; (iv) extreme loss, species with changes in range size between -0.8 and -0.99; and (v) complete loss, species with changes in range size of -1.0.

### Climatic characterization of ΔSR

We explored whether areas with declining or increasing species richness (SR) showed climatic differences in the present. For this, we used the resulting per-country maps of ΔSR to characterize areas with estimated positive and negative ΔSR (gains and losses, respectively) in terms of their bioclimatic profile. After estimating ΔSR for each country, we identified grid-cells with the largest gains and losses in species richness (positive and negative ΔSR) as those with values above the third quantiles and below the first quantile of the distribution of ΔSR, respectively. We characterized areas of loss and gains using four bioclimatic variables: mean annual temperature, temperature seasonality, total annual precipitation, and precipitation seasonality (**Supplementary Methods**).

We characterized the bioclimatic profile across countries in order to explore the influence of different variables (*e*.*g*., temperature, precipitation) on species responses. For this, we estimated the temporal change in four bioclimatic variables (Δbio) across countries under the different scenarios and temporal horizons (**Supplementary Methods**).

## Acknowledgements

We are grateful to Francisco Estrada-Porrúa, Julián A Velasco and the members of group “Clima y Sociedad” for the ideas that prompted the development of the project and comments on earlier drafts. Part of the analyses in this paper were carried out on CONABIO’s (Comisión Nacional para el Conocimiento y Uso de la Biodiversidad) computing cluster, supported by their system administrator and the Subcoordinación de soporte informático; and the other part of the analyses in this paper were carried out on the computing cluster Tláloc (Centro de Ciencias de la Atmósfera) supported by their system administrator.

## Author Contributions

C.U., A.P.C.-R. conceived the idea with the help of C.G.-G.; C.U., S.R.-B., and A.P.C.-R. designed the research and analyses; C.U. and A.P.C.-R. compiled and processed the data on animals; S.R.-B. compiled and processed the data on plants; D.S. and D.D. provided the climate simulations for melting scenarios; O.C.-B. generated and processed the bioclimate layers used in the modelling; P.C.-S. processed the data and performed the setup of the computing cluster for the analyses; C.U., S.R.-B., and A.P.C.-R. performed the analyses; C.U. and S.R.-B. lead the writing with contributions from A.P.C.-R.; C.U., S.R.-B., D.S., D.D. and A.P.C.-R. discussed the manuscript. All authors read and approved the manuscript.

## Data availability

Data for species distribution models are available at Zenodo with the identifier doi.org/10.5281/zenodo.4917258. The geographic occurrence data for vascular plants is available from the Global Biodiversity Information Facility with the identifier https://doi.org/10.15468/dl.bdxzkw. The distribution polygons for tetrapods and vascular plants are available at https://www.iucnredlist.org/ and http://www.birdlife.org/.

## Code availability

All R code used for processing the distribution models, and to perform the geospatial and statistical analyses are available at Zenodo with the identifier doi.org/10.5281/zenodo.4917258.

## Extended Data

**Extended Data Figure 1.**
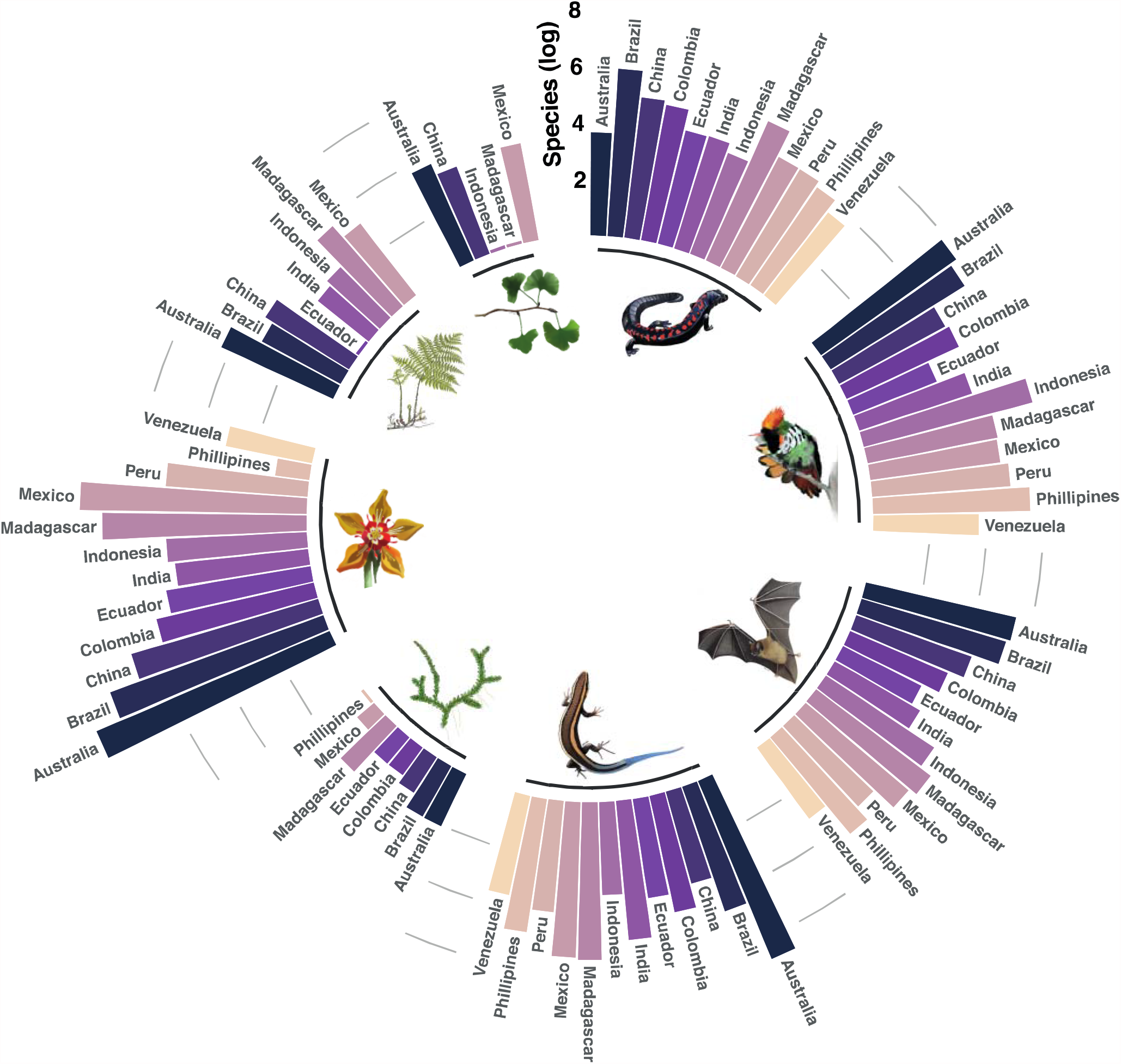
Number of species modeled for each group of vascular plants and tetrapods across the twelve megadiverse countries. Species numbers are given on a log scale. Groups of species are depicted by the inset illustrations (from top, clockwise): amphibians, birds, mammals, reptiles, lycophytes, flowering plants, ferns, and gymnosperms.

**Extended Data Figure 2.**
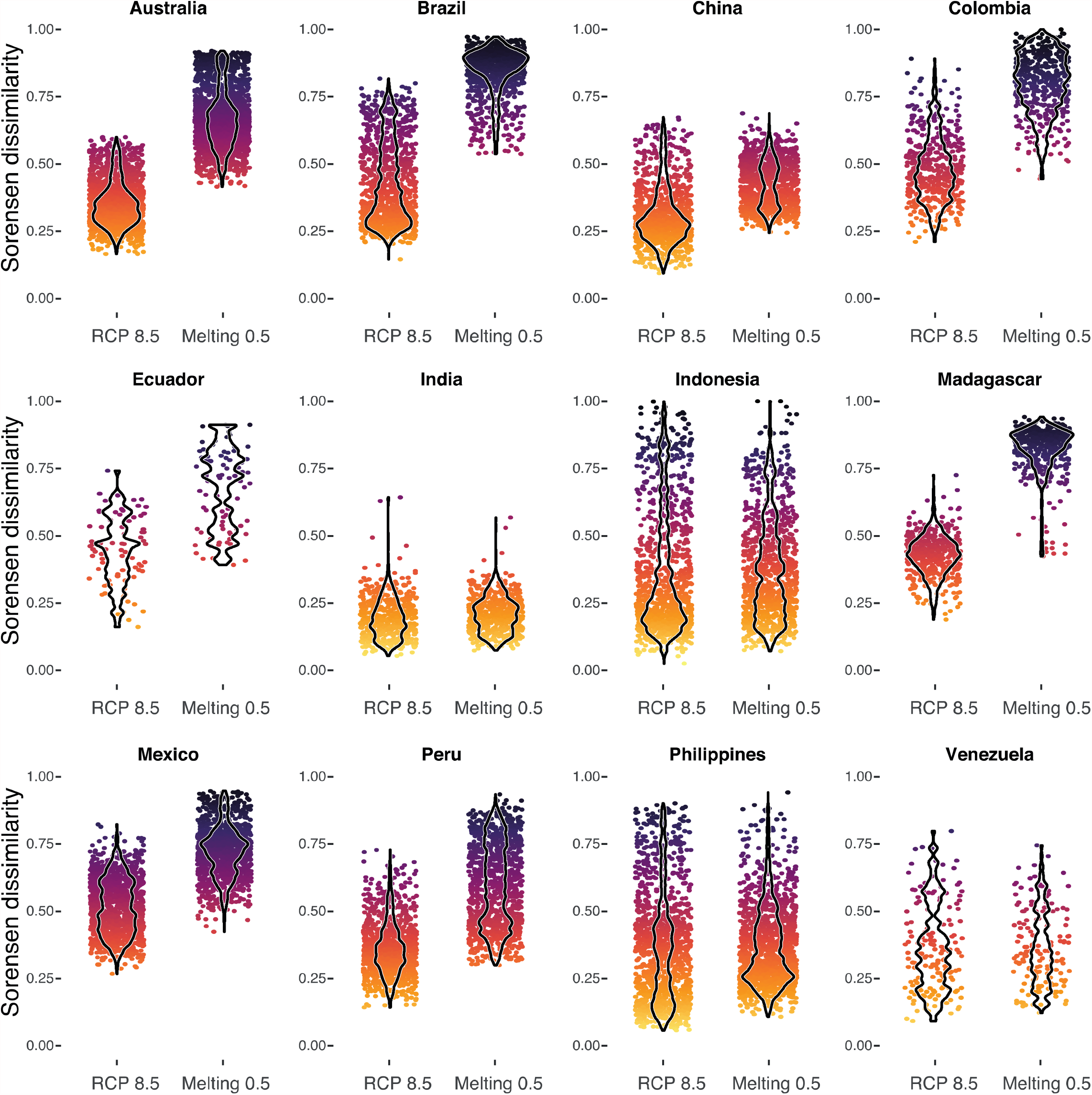
Changes in species composition within potential species hotspots (PSH) across the twelve megadiverse countries. Temporal changes in species composition were based on species distribution models (SDMs) and were estimated using the Sørensen dissimilarity index (β_SøR_) for individual pixels across time. Values approaching one indicate increasing dissimilarity in composition across time. Changes in composition are shown across countries for 2030 under the RCP 8.5 and Melting 0.5 scenarios.

**Extended Data Figure 3.**
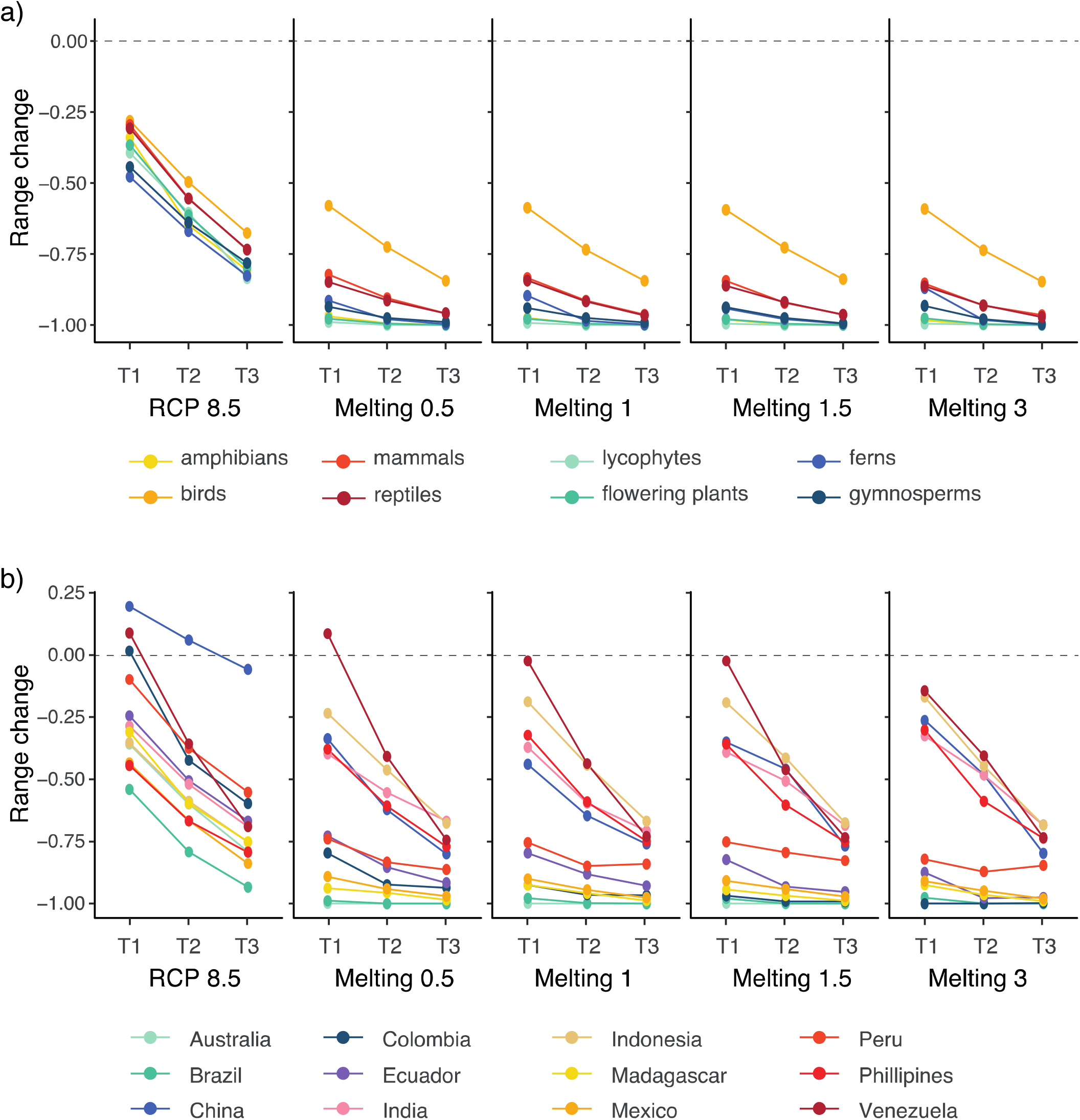
Changes in the size of the distribution range of species across taxonomic groups and countries under five scenarios of climate change at three different time horizons. a–b, estimated median range size change for the eight (a) taxonomic groups and (b) countries across scenarios and time horizons. Species range sizes were standardized relative to the present-day species range size and then summarized across groups: positive values indicate range reductions (1 means complete loss) and negative values indicate range expansions. T1: 2030; T2: 2050; T3: 2070.

**Extended Data Figure 4.**
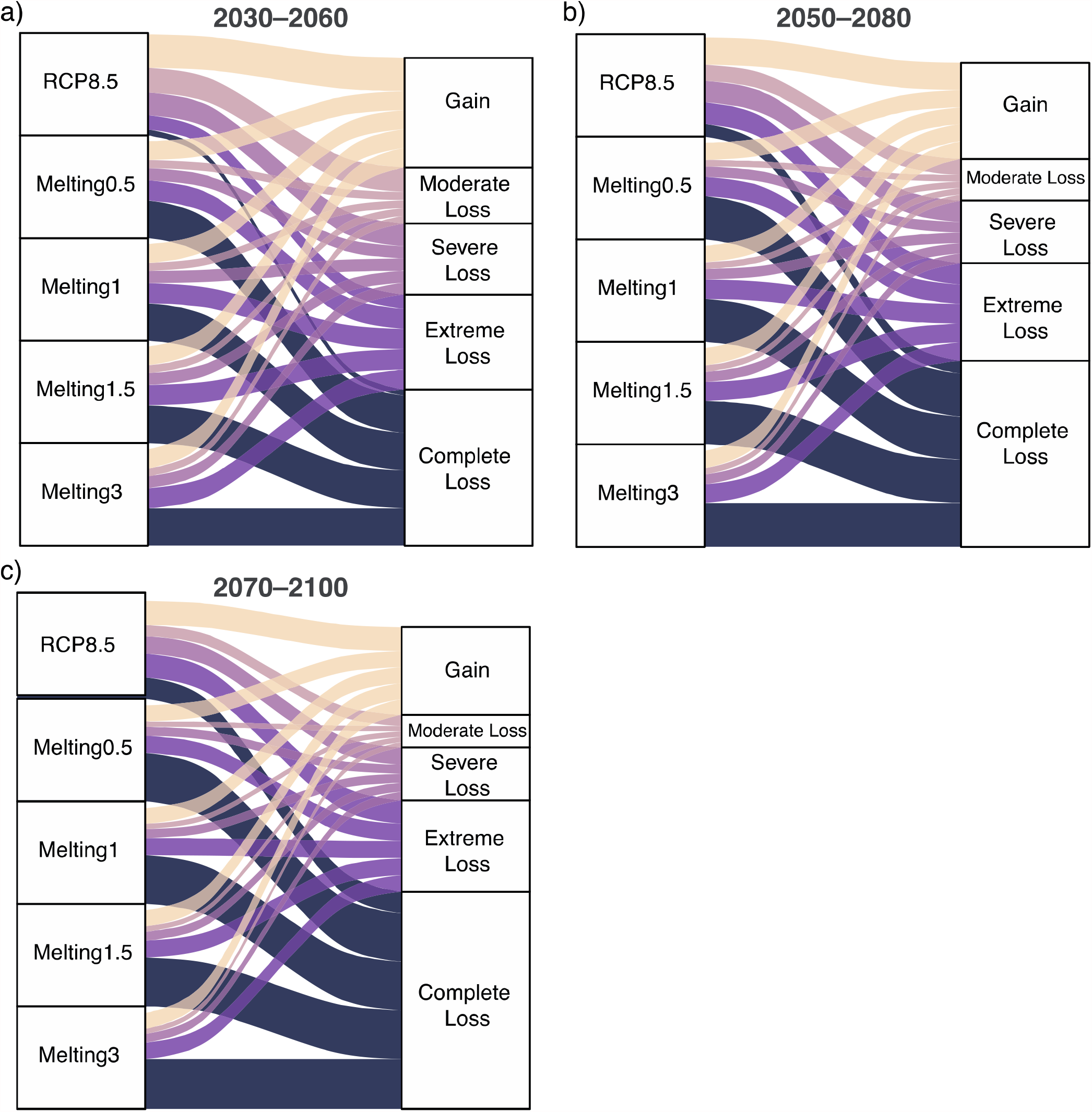
Proportion of species falling within different categories of change in species’ range sizes under different climate change scenarios. a-c, alluvial plots showing the distribution of five categories of change in species’ range size under the RCP 8.5 and the four Melting scenarios. The vertical size of the blocks and the width of the flows are proportional to the frequency of species within each block/flow. All scenarios have the same block size corresponding to the 21,146 modelled species. The flows represent the proportion of species within either of the five categories as estimated under each of the five scenarios. Range size estimation was based on species distribution models and were standardized relative to the present-day species range size.

