## Supplementary Information for "Greenland’s thaw pushes the biodiversity crisis"

• This file contains Supplementary Methods, Supplementary tables 1–6, Supplementary figures 1–9, and a list of supplementary data files.
• Supplementary data files and R code are available at Zenodo with the identifier http://doi.org/10.5281/zenodo.4917258
• Data and code citation (version 1.0):

Carolina Ureta, Santiago Ramírez Barahona, Óscar Calderón-Bustamante, Pedro Cruz-Santiago, Carlos Gay-García, Didier Swingedouw, Angela P. Cuervo-Robayo (2021)
*Greenland's thaw pushes the biodiversity crisis* (Version v1.0) [Data set]. Zenodo.
http://doi.org/10.5281/zenodo.4917258

### Supplementary Methods

#### Species selection and modelling

We selected a total of 21,252 species of vascular plants and tetrapods for species distribution modelling across twelve megadiverse countries (**Extended Data figure 1**). The selection of species was based on two criteria: (i) species endemic to any country; and (ii) species with more than 25 unique occurrence records. In sum, we modelled a total of 5,757 tetrapod species and 15,389 vascular plant species, with a broad representation of species for the eight groups across the world. The validation statistics (TSS, ROC) show an overall good performance of the ensemble models (**Supplementary figure 10**). Out of the 21,146 modelled species, 89% had ensemble models with $TSS \geq 0.7$, whereas 98% had ensemble models with $ROC \geq 0.85$.

#### Potential species hotspots

In our main analyses, we defined potential species hotspots (PSHs) using a threshold of 60% of the highest level of species richness observed in the present-day. Given that we use the threshold to define PSHs, their extent is expected to be strongly influenced by the choice of threshold. Therefore, we tested two additional thresholds (20 and 90%) to define and quantify the extent of PSHs. As expected, the extent of PSHs across megadiverse countries varies considerably when applying different thresholds. However, these additional results agree with the general trend we discuss in the main text; a drastic reduction in the extent of areas harboring substantial species richness (**Supplementary table S1**), and a less steep decline in geographic extent for areas with lower species richness (**figure 2** in main text).

### Species dissimilarity

We estimated the temporal species dissimilarity across PSHs using the Sørensen pairwise dissimilarity index ($\beta_{S\emptyset R}$). We implemented the estimation of $\beta_{S\emptyset R}$ for grid-cell $k$ between times $i$ and $j$ as

$$\beta_{ij}^k = (b + c) / (2 \times a + b + c)$$

where $a$ is the number of species in $k$ shared between times, $b$ are the species unique to time $i$, and $c$ are the species unique to time $j$. We estimated $\beta_{S\emptyset R}$ under each scenario and temporal horizon and show that PSHs are expected to undergo substantial temporal changes in species composition, specially under the melting scenarios (**Supplementary table S2**).

### Bioclimatic characterization

We wanted to assess whether the loss (or gain) in SR was partly influenced by climate. Thus, we characterized areas with estimated positive and negative $\Delta SR$ (gains and losses, respectively) in term of their bioclimatic profile. For this, we extracted data for four bioclimatic variables – at the present – across grid-cells and then classified grid-cells according to their $\Delta SR$. We estimated and visualized differences between areas of loss and gain across countries under two scenarios: RCPS 8.5 and Melting 0.5 (**Supplementary figures S3–S6**).

### Species Richness

To complement the data provided in the main text, we visualized SR and ΔSR for vascular plants and tetrapods separately, showing that the overall response of vascular plants and tetrapods varies by country (**Supplementary Figure S1–S2**). Given that the distribution of SR among the four melting scenarios is fairly stable, we only visualized differences in the spatial distribution of SR between the two groups under the RCP 8.5 and the Melting 0.5 scenarios. Species richness estimates – and the corresponding ΔSR – are available as supplementary data.

We quantified proportional changes in species' distribution ranges under each scenario and temporal horizon as

$$\frac{range_{present} - range_{future}}{range_{present}}$$

where ranges were estimated as the number of grid-cells occupied by species. Positive values indicate range contractions, whereas negative values indicate range expansions. We summarized changes in species' ranges across groups and countries using density distributions (**figure 3** in main text) and point estimates (**Supplementary tables S3–S4**). We used the per-species estimates to quantify the proportion of species with a complete loss of distribution area (that is, potential extirpation). We visualized differences between the RCP 8.5 scenario and the four melting scenarios across groups and countries (**Supplementary figures S7–S8**); these data are summarized in **Supplementary tables S5–S6**. These data support our main findings of an enhanced impact of melting on species persistence across megadiverse countries, but that differences among melting scenarios are small.

### Supplementary tables and figures

#### Supplementary table S1

**Projected extent of potential species hotspots across twelve megadiverse countries.** Values were estimated for each country and represent the geographic extent (relative to their present) that is projected under five climate change scenarios and three temporal horizons.

| Scenario | Australia | Brazil | China | Colombia | Ecuador | India |
| --- | --- | --- | --- | --- | --- | --- |
| *2020–2060* |  |  |  |  |  |  |
| RCP 8.5 | 0.689 | 0.090 | 1.527 | 0.882 | 0.150 | 0.470 |
| Scenario 0.5 | 0.000 | 0.000 | 2.733 | 0.003 | 0.000 | 0.608 |
| Scenario 1 | 0.000 | 0.000 | 2.184 | 0.000 | 0.000 | 0.628 |
| Scenario 1.5 | 0.000 | 0.000 | 2.737 | 0.000 | 0.000 | 0.606 |
| Scenario 3 | 0.000 | 0.000 | 2.642 | 0.000 | 0.000 | 0.616 |
| *2050–2080* |  |  |  |  |  |  |
| RCP 8.5 | 0.302 | 0.007 | 2.068 | 0.093 | 0.163 | 0.308 |
| Scenario 0.5 | 0.000 | 0.000 | 0.744 | 0.000 | 0.000 | 0.454 |
| Scenario 1 | 0.000 | 0.000 | 0.662 | 0.000 | 0.000 | 0.428 |
| Scenario 1.5 | 0.000 | 0.000 | 2.113 | 0.000 | 0.000 | 0.494 |
| Scenario 3 | 0.000 | 0.000 | 1.526 | 0.000 | 0.000 | 0.481 |
| *2070–2100* |  |  |  |  |  |  |
| RCP 8.5 | 0.066 | 0.000 | 2.497 | 0.119 | 0.000 | 0.209 |
| Scenario 0.5 | 0.000 | 0.000 | 0.245 | 0.000 | 0.000 | 0.315 |
| Scenario 1 | 0.000 | 0.000 | 0.238 | 0.000 | 0.000 | 0.286 |
| Scenario 1.5 | 0.000 | 0.000 | 0.375 | 0.000 | 0.000 | 0.293 |
| Scenario 3 | 0.000 | 0.000 | 0.177 | 0.000 | 0.000 | 0.339 |

**Supplementary table S1. Continued.**

| Scenario | Indonesia | Madagascar | Mexico | Peru | Phillippines | Venezuela |
| --- | --- | --- | --- | --- | --- | --- |
| *2020–2060* |  |  |  |  |  |  |
| RCP 8.5 | 0.521 | 0.199 | 0.540 | 0.883 | 0.351 | 1.399 |
| Scenario 0.5 | 0.304 | 0.000 | 0.015 | 0.255 | 0.401 | 1.266 |
| Scenario 1 | 0.297 | 0.000 | 0.006 | 0.256 | 0.450 | 1.374 |
| Scenario 1.5 | 0.321 | 0.000 | 0.014 | 0.262 | 0.457 | 1.247 |
| Scenario 3 | 0.344 | 0.000 | 0.011 | 0.240 | 0.534 | 1.076 |
| *2050–2080* |  |  |  |  |  |  |
| RCP 8.5 | 0.243 | 0.015 | 0.263 | 0.638 | 0.157 | 0.507 |
| Scenario 0.5 | 0.166 | 0.000 | 0.006 | 0.200 | 0.199 | 0.564 |
| Scenario 1 | 0.166 | 0.000 | 0.000 | 0.300 | 0.199 | 0.475 |
| Scenario 1.5 | 0.146 | 0.000 | 0.001 | 0.314 | 0.204 | 0.418 |
| Scenario 3 | 0.182 | 0.000 | 0.000 | 0.285 | 0.239 | 0.583 |
| *2070–2100* |  |  |  |  |  |  |
| RCP 8.5 | 0.147 | 0.000 | 0.085 | 0.392 | 0.083 | 0.152 |
| Scenario 0.5 | 0.095 | 0.000 | 0.000 | 0.247 | 0.111 | 0.133 |
| Scenario 1 | 0.054 | 0.000 | 0.000 | 0.329 | 0.112 | 0.089 |
| Scenario 1.5 | 0.063 | 0.000 | 0.000 | 0.303 | 0.115 | 0.140 |
| Scenario 3 | 0.051 | 0.000 | 0.000 | 0.393 | 0.115 | 0.140 |

### Supplementary table S2

**Projected temporal dissimilarity in species composition across twelve megadiverse countries.** Values were estimated using the Sørensen dissimilarity index ($\beta_{S\text{ØR}}$) and represent the median dissimilarity between present-day species composition and the future projected composition under five climate change scenarios and three temporal horizons.

| Scenario | Australia | Brazil | China | Colombia | Ecuador | India |
| --- | --- | --- | --- | --- | --- | --- |
| *2020–2060* |  |  |  |  |  |  |
| RCP 8.5 | 0.351 | 0.429 | 0.305 | 0.481 | 0.443 | 0.192 |
| Scenario 0.5 | 0.667 | 0.853 | 0.426 | 0.801 | 0.677 | 0.208 |
| Scenario 1 | 0.664 | 0.849 | 0.442 | 0.869 | 0.712 | 0.196 |
| Scenario 1.5 | 0.656 | 0.854 | 0.436 | 0.883 | 0.740 | 0.208 |
| Scenario 3 | 0.638 | 0.829 | 0.456 | 0.916 | 0.717 | 0.213 |
| *2050–2080* |  |  |  |  |  |  |
| RCP 8.5 | 0.483 | 0.574 | 0.358 | 0.731 | 0.603 | 0.330 |
| Scenario 0.5 | 0.706 | 0.932 | 0.498 | 0.889 | 0.747 | 0.250 |
| Scenario 1 | 0.709 | 0.924 | 0.501 | 0.913 | 0.768 | 0.271 |
| Scenario 1.5 | 0.702 | 0.932 | 0.468 | 0.916 | 0.783 | 0.240 |
| Scenario 3 | 0.695 | 0.929 | 0.473 | 0.933 | 0.752 | 0.259 |
| *2070–2100* |  |  |  |  |  |  |
| RCP 8.5 | 0.607 | 0.687 | 0.423 | 0.858 | 0.790 | 0.433 |
| Scenario 0.5 | 0.747 | 0.961 | 0.555 | 0.913 | 0.778 | 0.289 |
| Scenario 1 | 0.762 | 0.960 | 0.568 | 0.936 | 0.777 | 0.309 |
| Scenario 1.5 | 0.756 | 0.958 | 0.563 | 0.939 | 0.799 | 0.302 |
| Scenario 3 | 0.769 | 0.960 | 0.569 | 0.946 | 0.804 | 0.291 |

**Supplementary table S2. Continued.**

| Scenario | Indonesia | Madagascar | Mexico | Peru | Philippines | Venezuela |
| --- | --- | --- | --- | --- | --- | --- |
| *2020–2060* |  |  |  |  |  |  |
| RCP 8.5 | 0.362 | 0.426 | 0.516 | 0.363 | 0.364 | 0.363 |
| Scenario 0.5 | 0.374 | 0.824 | 0.720 | 0.593 | 0.370 | 0.362 |
| Scenario 1 | 0.406 | 0.808 | 0.716 | 0.573 | 0.356 | 0.361 |
| Scenario 1.5 | 0.346 | 0.818 | 0.735 | 0.573 | 0.366 | 0.365 |
| Scenario 3 | 0.338 | 0.806 | 0.722 | 0.603 | 0.348 | 0.396 |
| *2050–2080* |  |  |  |  |  |  |
| RCP 8.5 | 0.554 | 0.617 | 0.655 | 0.485 | 0.506 | 0.502 |
| Scenario 0.5 | 0.541 | 0.854 | 0.785 | 0.636 | 0.472 | 0.428 |
| Scenario 1 | 0.498 | 0.860 | 0.779 | 0.614 | 0.473 | 0.421 |
| Scenario 1.5 | 0.513 | 0.865 | 0.781 | 0.609 | 0.474 | 0.428 |
| Scenario 3 | 0.509 | 0.864 | 0.780 | 0.631 | 0.474 | 0.433 |
| *2070–2100* |  |  |  |  |  |  |
| RCP 8.5 | 0.665 | 0.727 | 0.765 | 0.607 | 0.609 | 0.649 |
| Scenario 0.5 | 0.651 | 0.890 | 0.822 | 0.659 | 0.572 | 0.521 |
| Scenario 1 | 0.659 | 0.892 | 0.829 | 0.658 | 0.565 | 0.513 |
| Scenario 1.5 | 0.680 | 0.892 | 0.825 | 0.666 | 0.568 | 0.501 |
| Scenario 3 | 0.677 | 0.897 | 0.832 | 0.650 | 0.565 | 0.514 |

#### Supplementary table S3

**Projected range loss for vascular plant and tetrapod species across twelve megadiverse countries.** Values were estimated for each group and represent the median range loss of species (relative to their present extent) that is projected under five climate change scenarios and three temporal horizons.

| Scenario | Lycophytes | Flowering plants | Ferns | Gymnosperms |
| --- | --- | --- | --- | --- |
| *2020–2060* |  |  |  |  |
| RCP 8.5 | -0.393 | -0.366 | -0.477 | -0.442 |
| Scenario 0.5 | -0.989 | -0.977 | -0.914 | -0.935 |
| Scenario 1 | -0.992 | -0.978 | -0.896 | -0.939 |
| Scenario 1.5 | -0.995 | -0.978 | -0.940 | -0.937 |
| Scenario 3 | -0.995 | -0.976 | -0.869 | -0.931 |
| *2050–2080* |  |  |  |  |
| RCP 8.5 | -0.604 | -0.613 | -0.668 | -0.638 |
| Scenario 0.5 | -1.000 | -0.995 | -0.979 | -0.975 |
| Scenario 1 | -1.000 | -0.995 | -0.984 | -0.974 |
| Scenario 1.5 | -1.000 | -0.995 | -0.979 | -0.974 |
| Scenario 3 | -0.998 | -0.996 | -0.982 | -0.978 |
| *2070–2100* |  |  |  |  |
| RCP 8.5 | -0.834 | -0.797 | -0.826 | -0.781 |
| Scenario 0.5 | -1.000 | -1.000 | -0.996 | -0.989 |
| Scenario 1 | -1.000 | -1.000 | -0.998 | -0.991 |
| Scenario 1.5 | -1.000 | -1.000 | -0.994 | -0.994 |
| Scenario 3 | -1.000 | -1.000 | -0.998 | -0.996 |

**Supplementary table S3. Continued.**

| Scenario | Amphibians | Birds | Mammals | Reptiles |
| --- | --- | --- | --- | --- |
| *2020–2060* |  |  |  |  |
| RCP 8.5 | -0.339 | -0.280 | -0.294 | -0.306 |
| Scenario 0.5 | -0.969 | -0.579 | -0.821 | -0.848 |
| Scenario 1 | -0.975 | -0.586 | -0.833 | -0.843 |
| Scenario 1.5 | -0.980 | -0.593 | -0.843 | -0.861 |
| Scenario 3 | -0.983 | -0.590 | -0.853 | -0.863 |
| *2050–2080* |  |  |  |  |
| RCP 8.5 | -0.648 | -0.495 | -0.553 | -0.555 |
| Scenario 0.5 | -0.994 | -0.725 | -0.905 | -0.912 |
| Scenario 1 | -0.996 | -0.734 | -0.913 | -0.916 |
| Scenario 1.5 | -0.998 | -0.727 | -0.921 | -0.918 |
| Scenario 3 | -0.997 | -0.736 | -0.931 | -0.930 |
| *2070–2100* |  |  |  |  |
| RCP 8.5 | -0.806 | -0.675 | -0.734 | -0.733 |
| Scenario 0.5 | -1.000 | -0.843 | -0.958 | -0.957 |
| Scenario 1 | -1.000 | -0.844 | -0.962 | -0.965 |
| Scenario 1.5 | -1.000 | -0.838 | -0.962 | -0.962 |
| Scenario 3 | -1.000 | -0.847 | -0.964 | -0.971 |

#### Supplementary table S4

**Projected range loss for vascular plant and tetrapod species across twelve megadiverse countries.** Values were estimated for each country and represent the median range loss of species (relative to their present extent) that is projected under five climate change scenarios and three temporal horizons.

| Scenario | Australia | Brazil | China | Colombia | Ecuador | India |
| --- | --- | --- | --- | --- | --- | --- |
| *2020–2060* |  |  |  |  |  |  |
| RCP 8.5 | -0.358 | -0.540 | 0.196 | 0.017 | -0.244 | -0.285 |
| Scenario 0.5 | -0.999 | -0.988 | -0.336 | -0.796 | -0.727 | -0.397 |
| Scenario 1 | -0.999 | -0.978 | -0.439 | -0.924 | -0.796 | -0.371 |
| Scenario 1.5 | -0.999 | -0.979 | -0.349 | -0.968 | -0.823 | -0.390 |
| Scenario 3 | -0.998 | -0.975 | -0.262 | -1.000 | -0.874 | -0.325 |
| *2050–2080* |  |  |  |  |  |  |
| RCP 8.5 | -0.598 | -0.792 | 0.061 | -0.423 | -0.504 | -0.519 |
| Scenario 0.5 | -1.000 | -1.000 | -0.622 | -0.923 | -0.853 | -0.553 |
| Scenario 1 | -1.000 | -0.998 | -0.646 | -0.964 | -0.880 | -0.595 |
| Scenario 1.5 | -1.000 | -0.999 | -0.458 | -0.991 | -0.932 | -0.506 |
| Scenario 3 | -1.000 | -0.998 | -0.480 | -1.000 | -0.977 | -0.483 |
| *2070–2100* |  |  |  |  |  |  |
| RCP 8.5 | -0.789 | -0.933 | -0.058 | -0.597 | -0.667 | -0.690 |
| Scenario 0.5 | -1.000 | -1.000 | -0.800 | -0.935 | -0.916 | -0.667 |
| Scenario 1 | -1.000 | -1.000 | -0.759 | -0.966 | -0.928 | -0.707 |
| Scenario 1.5 | -1.000 | -1.000 | -0.768 | -0.991 | -0.952 | -0.685 |
| Scenario 3 | -1.000 | -1.000 | -0.797 | -0.997 | -0.974 | -0.683 |

**Supplementary table S4. Continued.**

| Scenario | Indonesia | Madagascar | Mexico | Peru | Philippines | Venezuela |
| --- | --- | --- | --- | --- | --- | --- |
| *2020–2060* |  |  |  |  |  |  |
| RCP 8.5 | -0.352 | -0.309 | -0.434 | -0.098 | -0.444 | 0.089 |
| Scenario 0.5 | -0.234 | -0.938 | -0.891 | -0.739 | -0.378 | 0.087 |
| Scenario 1 | -0.187 | -0.925 | -0.900 | -0.754 | -0.321 | -0.023 |
| Scenario 1.5 | -0.191 | -0.943 | -0.907 | -0.751 | -0.357 | -0.023 |
| Scenario 3 | -0.169 | -0.924 | -0.909 | -0.821 | -0.301 | -0.143 |
| *2050–2080* |  |  |  |  |  |  |
| RCP 8.5 | -0.588 | -0.597 | -0.666 | -0.375 | -0.667 | -0.357 |
| Scenario 0.5 | -0.462 | -0.956 | -0.940 | -0.833 | -0.607 | -0.408 |
| Scenario 1 | -0.442 | -0.958 | -0.944 | -0.848 | -0.590 | -0.436 |
| Scenario 1.5 | -0.414 | -0.967 | -0.941 | -0.793 | -0.603 | -0.461 |
| Scenario 3 | -0.446 | -0.964 | -0.948 | -0.870 | -0.588 | -0.405 |
| *2070–2100* |  |  |  |  |  |  |
| RCP 8.5 | -0.752 | -0.750 | -0.838 | -0.551 | -0.792 | -0.689 |
| Scenario 0.5 | -0.675 | -0.985 | -0.969 | -0.862 | -0.770 | -0.743 |
| Scenario 1 | -0.668 | -0.987 | -0.975 | -0.840 | -0.748 | -0.730 |
| Scenario 1.5 | -0.674 | -0.988 | -0.972 | -0.826 | -0.751 | -0.734 |
| Scenario 3 | -0.684 | -0.989 | -0.980 | -0.846 | -0.736 | -0.734 |

#### Supplementary table S5

**Proportion of vascular plant and tetrapod species projected with complete range loss across twelve megadiverse countries.** Values were estimated for each group and represent the proportion of species (relative to the total number of modeled species per country) with complete range reduction projected under five climate change scenarios and three temporal horizons.

| Scenario | Lycophytes | Flowering plants | Ferns | Gymnosperms |
| --- | --- | --- | --- | --- |
| *2020–2060* |  |  |  |  |
| RCP 8.5 | 0.086 | 0.061 | 0.025 | 0.044 |
| Scenario 0.5 | 0.400 | 0.406 | 0.365 | 0.359 |
| Scenario 1 | 0.429 | 0.402 | 0.350 | 0.337 |
| Scenario 1.5 | 0.400 | 0.405 | 0.365 | 0.370 |
| Scenario 3 | 0.400 | 0.397 | 0.350 | 0.359 |
| *2050–2080* |  |  |  |  |
| RCP 8.5 | 0.115 | 0.131 | 0.085 | 0.077 |
| Scenario 0.5 | 0.515 | 0.462 | 0.425 | 0.414 |
| Scenario 1 | 0.515 | 0.457 | 0.425 | 0.392 |
| Scenario 1.5 | 0.515 | 0.463 | 0.400 | 0.392 |
| Scenario 3 | 0.486 | 0.467 | 0.395 | 0.370 |
| *2070–2100* |  |  |  |  |
| RCP 8.5 | 0.229 | 0.223 | 0.170 | 0.196 |
| Scenario 0.5 | 0.572 | 0.516 | 0.470 | 0.457 |
| Scenario 1 | 0.515 | 0.457 | 0.425 | 0.392 |
| Scenario 1.5 | 0.572 | 0.517 | 0.465 | 0.446 |
| Scenario 3 | 0.600 | 0.530 | 0.475 | 0.446 |

**Supplementary table S5. Continued.**

| Scenario | Amphibians | Birds | Mammals | Reptiles |
| --- | --- | --- | --- | --- |
| *2020–2060* |  |  |  |  |
| RCP 8.5 | 0.074 | 0.040 | 0.061 | 0.056 |
| Scenario 0.5 | 0.403 | 0.162 | 0.270 | 0.268 |
| Scenario 1 | 0.414 | 0.158 | 0.270 | 0.266 |
| Scenario 1.5 | 0.430 | 0.160 | 0.270 | 0.277 |
| Scenario 3 | 0.445 | 0.171 | 0.277 | 0.283 |
| *2050–2080* |  |  |  |  |
| RCP 8.5 | 0.151 | 0.074 | 0.124 | 0.114 |
| Scenario 0.5 | 0.472 | 0.204 | 0.313 | 0.314 |
| Scenario 1 | 0.479 | 0.195 | 0.313 | 0.325 |
| Scenario 1.5 | 0.492 | 0.203 | 0.318 | 0.326 |
| Scenario 3 | 0.487 | 0.199 | 0.326 | 0.337 |
| *2070–2100* |  |  |  |  |
| RCP 8.5 | 0.235 | 0.123 | 0.197 | 0.196 |
| Scenario 0.5 | 0.526 | 0.243 | 0.358 | 0.362 |
| Scenario 1 | 0.479 | 0.195 | 0.313 | 0.325 |
| Scenario 1.5 | 0.545 | 0.248 | 0.363 | 0.377 |
| Scenario 3 | 0.526 | 0.249 | 0.378 | 0.381 |

### Supplementary table S6

**Proportion of vascular plant and tetrapod species projected with complete range loss across twelve megadiverse countries.** Values were estimated for each country and represent the proportion of species (relative to the total number of modeled species per country) with complete range reduction projected under five climate change scenarios and three temporal horizons.

| Scenario | Australia | Brazil | China | Colombia | Ecuador | India |
| --- | --- | --- | --- | --- | --- | --- |
| *2020–2060* |  |  |  |  |  |  |
| RCP 8.5 | 0.070 | 0.077 | 0.026 | 0.029 | 0.084 | 0.032 |
| Scenario 0.5 | 0.491 | 0.418 | 0.273 | 0.234 | 0.263 | 0.123 |
| Scenario 1 | 0.493 | 0.375 | 0.283 | 0.321 | 0.252 | 0.125 |
| Scenario 1.5 | 0.491 | 0.380 | 0.277 | 0.397 | 0.280 | 0.131 |
| Scenario 3 | 0.485 | 0.359 | 0.272 | 0.503 | 0.322 | 0.125 |
| *2050–2080* |  |  |  |  |  |  |
| RCP 8.5 | 0.146 | 0.192 | 0.050 | 0.055 | 0.161 | 0.075 |
| Scenario 0.5 | 0.535 | 0.513 | 0.309 | 0.301 | 0.305 | 0.157 |
| Scenario 1 | 0.536 | 0.474 | 0.319 | 0.348 | 0.333 | 0.147 |
| Scenario 1.5 | 0.536 | 0.484 | 0.299 | 0.463 | 0.368 | 0.149 |
| Scenario 3 | 0.536 | 0.485 | 0.295 | 0.527 | 0.431 | 0.147 |
| *2070–2100* |  |  |  |  |  |  |
| RCP 8.5 | 0.240 | 0.325 | 0.081 | 0.082 | 0.242 | 0.128 |
| Scenario 0.5 | 0.579 | 0.593 | 0.344 | 0.321 | 0.382 | 0.176 |
| Scenario 1 | 0.536 | 0.474 | 0.319 | 0.348 | 0.333 | 0.147 |
| Scenario 1.5 | 0.578 | 0.590 | 0.348 | 0.449 | 0.399 | 0.168 |
| Scenario 3 | 0.586 | 0.588 | 0.342 | 0.483 | 0.438 | 0.165 |

**Supplementary table S6. Continued.**

| Scenario | Indonesia | Madagascar | Mexico | Peru | Philippines | Venezuela |
| --- | --- | --- | --- | --- | --- | --- |
| *2020–2060* |  |  |  |  |  |  |
| RCP 8.5 | 0.064 | 0.028 | 0.053 | 0.039 | 0.022 | 0.063 |
| Scenario 0.5 | 0.112 | 0.318 | 0.233 | 0.213 | 0.106 | 0.152 |
| Scenario 1 | 0.103 | 0.301 | 0.250 | 0.237 | 0.104 | 0.159 |
| Scenario 1.5 | 0.093 | 0.324 | 0.252 | 0.235 | 0.104 | 0.173 |
| Scenario 3 | 0.103 | 0.292 | 0.260 | 0.300 | 0.098 | 0.187 |
| *2050–2080* |  |  |  |  |  |  |
| RCP 8.5 | 0.120 | 0.061 | 0.099 | 0.078 | 0.053 | 0.097 |
| Scenario 0.5 | 0.143 | 0.371 | 0.291 | 0.254 | 0.151 | 0.194 |
| Scenario 1 | 0.137 | 0.370 | 0.297 | 0.257 | 0.147 | 0.207 |
| Scenario 1.5 | 0.133 | 0.392 | 0.294 | 0.264 | 0.147 | 0.221 |
| Scenario 3 | 0.134 | 0.376 | 0.302 | 0.307 | 0.149 | 0.249 |
| *2070–2100* |  |  |  |  |  |  |
| RCP 8.5 | 0.193 | 0.134 | 0.164 | 0.141 | 0.113 | 0.166 |
| Scenario 0.5 | 0.192 | 0.429 | 0.344 | 0.283 | 0.193 | 0.228 |
| Scenario 1 | 0.137 | 0.370 | 0.297 | 0.257 | 0.147 | 0.207 |
| Scenario 1.5 | 0.192 | 0.438 | 0.346 | 0.293 | 0.183 | 0.242 |
| Scenario 3 | 0.189 | 0.442 | 0.376 | 0.298 | 0.191 | 0.256 |

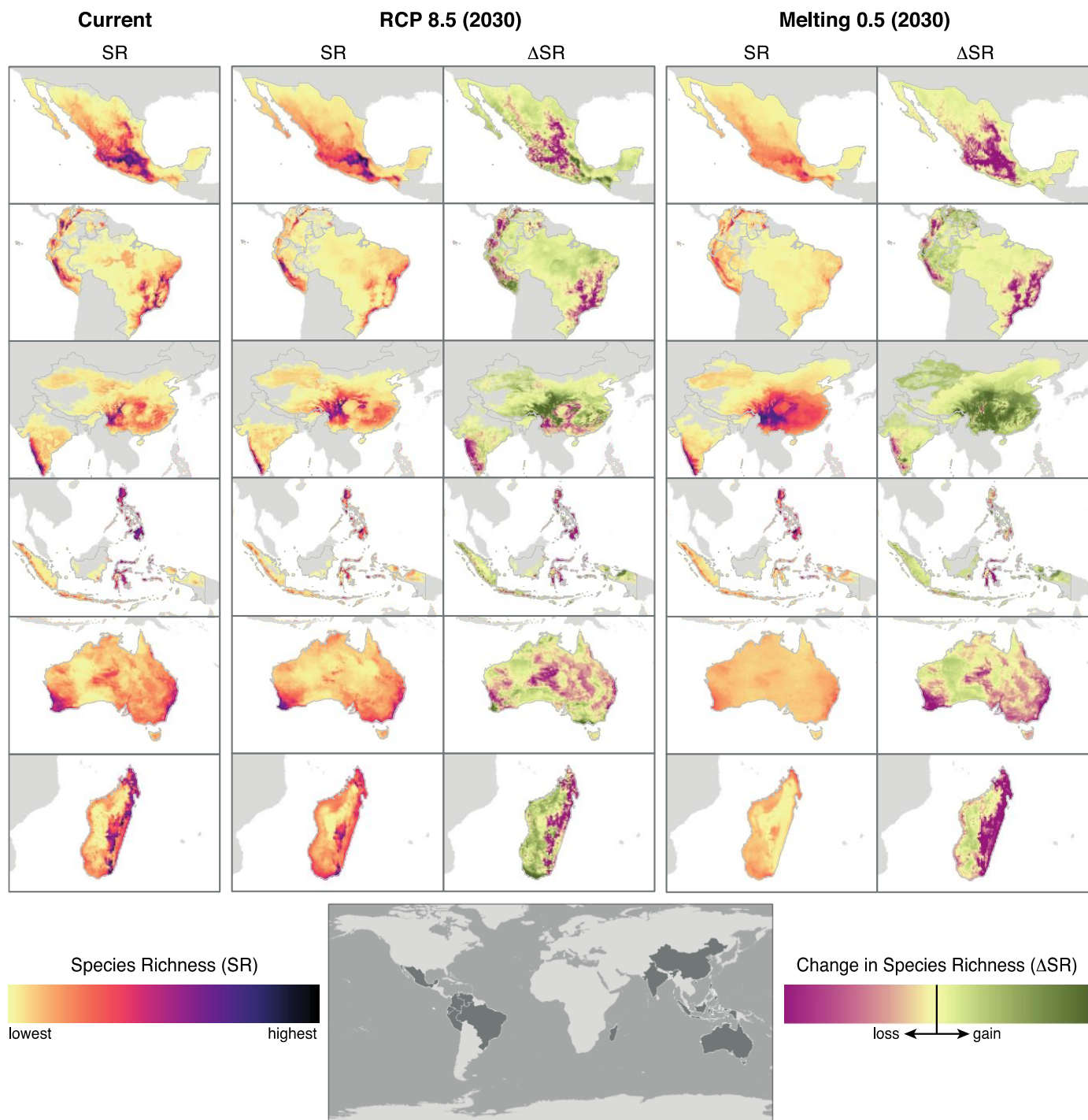

**Supplementary figure S1**

**Geographic patterns of present-day species richness and temporal changes in species richness of vascular plants across twelve megadiverse countries.** Estimates are based on species distribution models (SDMs) of vascular plants and tetrapods. Species richness (SR) was standardized to the range 0–1 within each country. Change in species richness (DSR) are shown for 2030–2060 under the RCP 8.5 and Melting 0.5 scenarios. For each grid-cell, the change in species richness was estimated as the difference in species richness estimated for the present-day and future scenarios.

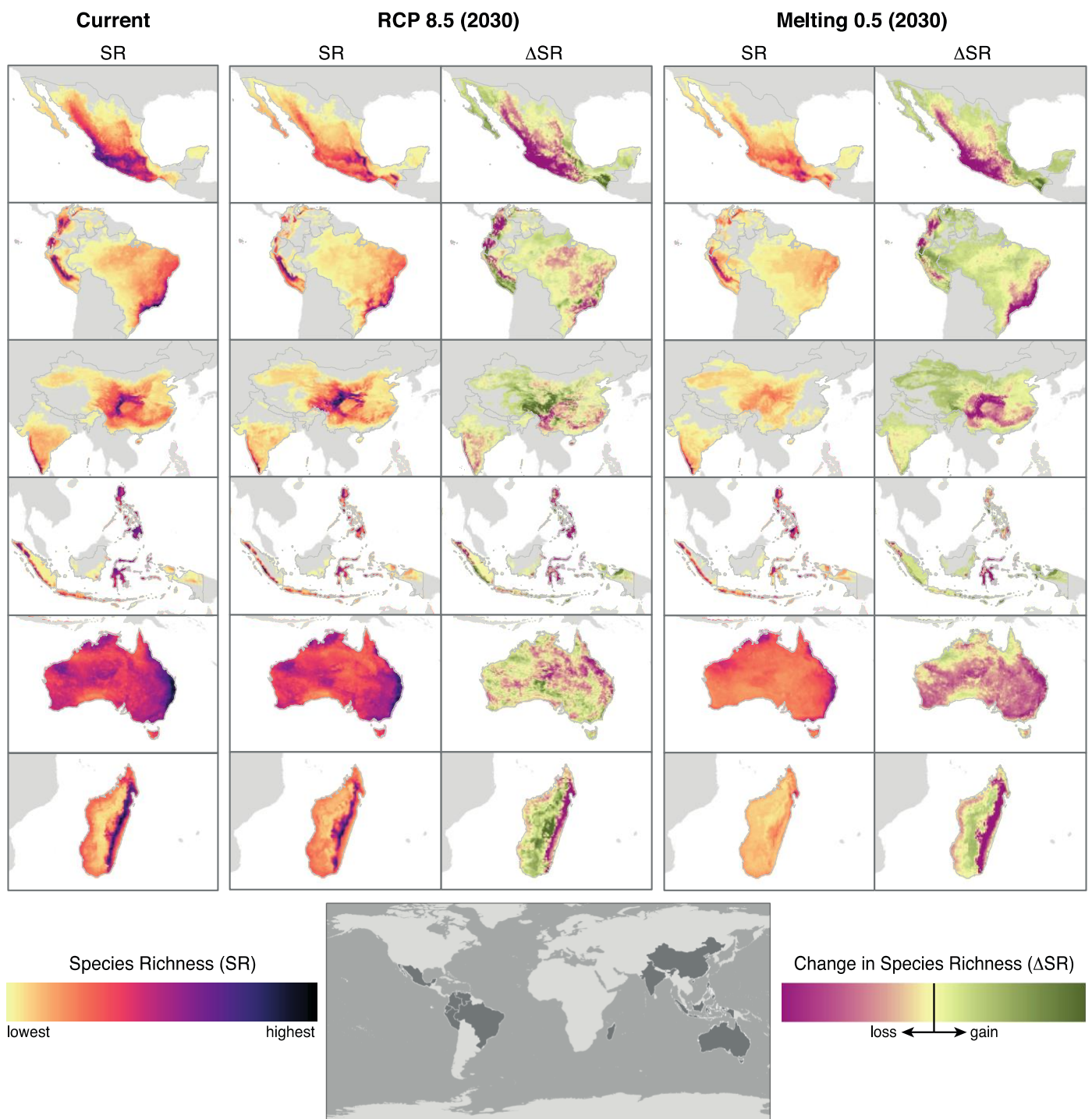

**Supplementary figure S2**

**Geographic patterns of present-day species richness and temporal changes in species richness of tetrapods across twelve megadiverse countries.** Estimates are based on species distribution models (SDMs) of vascular plants and tetrapods. Species richness (SR) was standardized to the range 0–1 within each country. Change in species richness (DSR) are shown for 2030–2060 under the RCP 8.5 and Melting 0.5 scenarios. For each grid-cell, the change in species richness was estimated as the difference in species richness estimated for the present-day and future scenarios.

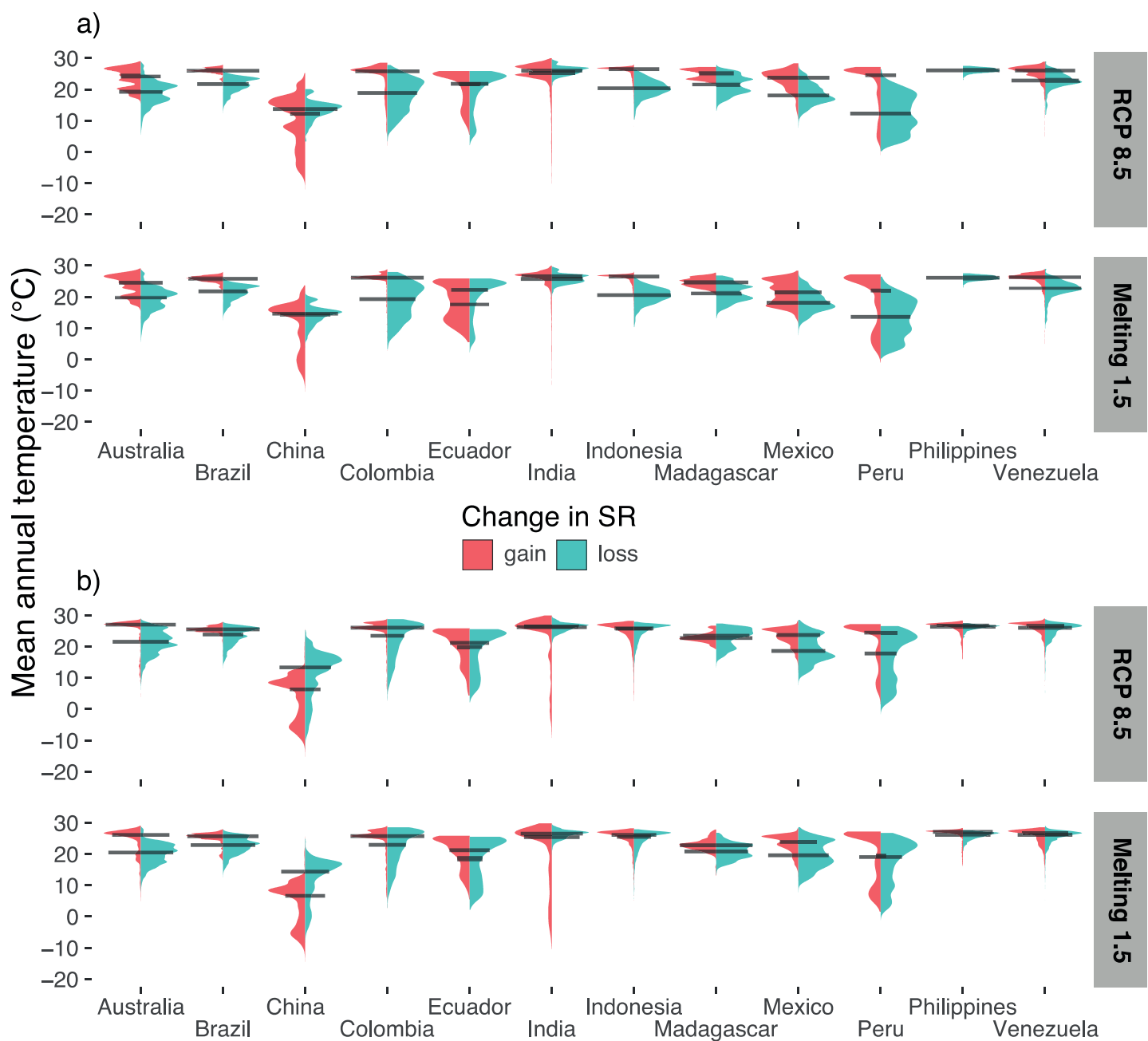

**Supplementary figure S3**

**Present-day mean annual temperature for areas projected with gain and loss in species richness across twelve megadiverse countries.** a–b, estimates are based on species distribution models (SDMs) of (a) vascular plants and (b) tetrapods. Areas of gain and loss represent pixels with ΔSR above the third quartile and below the first quartile, respectively. The bioclimatic profile for areas is show for 2030–2060 under the RCP 8.5 and Melting 1.5 scenarios.

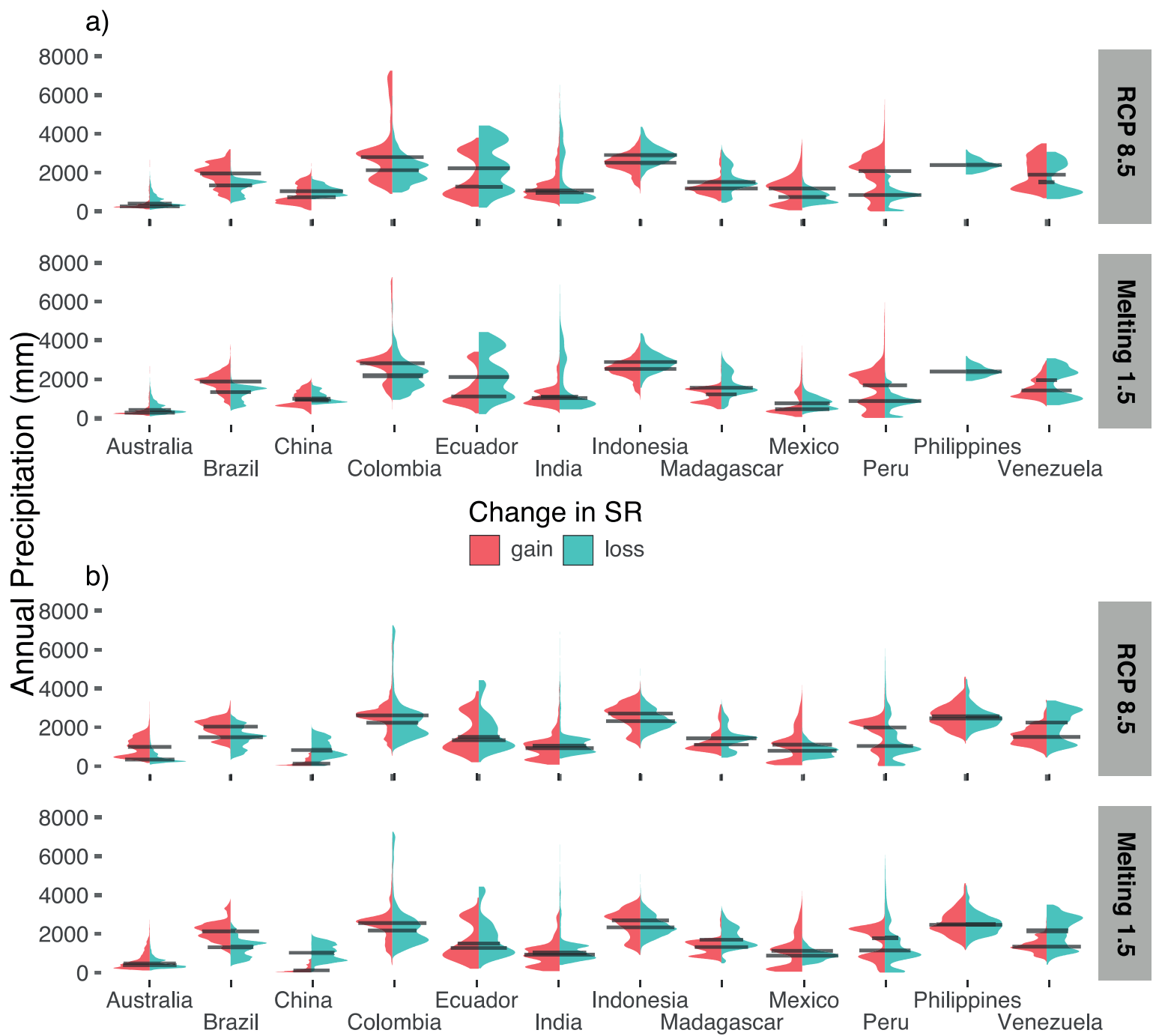

**Supplementary figure S4**

**Present-day annual precipitation for areas projected with gain and loss in species richness across twelve megadiverse countries.** a–b, estimates are based on species distribution models (SDMs) of (a) vascular plants and (b) tetrapods. Areas of gain and loss represent pixels with ΔSR above the third quartile and below the first quartile, respectively. The bioclimatic profile for areas is show for 2030–2060 under the RCP 8.5 and Melting 1.5 scenarios.

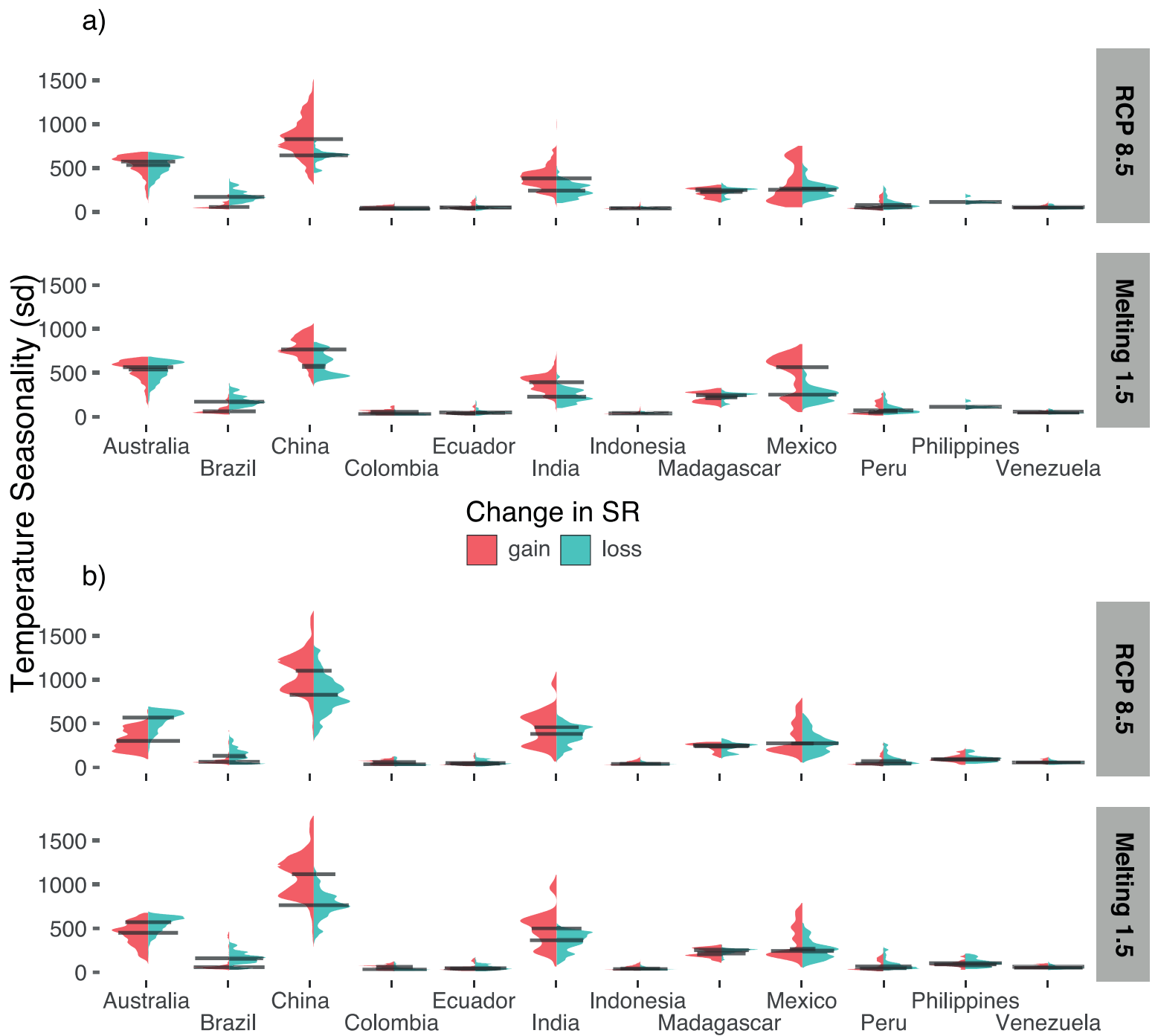

**Supplementary figure S5**

**Present-day temperature seasonality for areas projected with gain and loss in species richness across twelve megadiverse countries.** a–b, estimates are based on species distribution models (SDMs) of (a) vascular plants and (b) tetrapods. Areas of gain and loss represent pixels with ΔSR above the third quartile and below the first quartile, respectively. The bioclimatic profile for areas is show for 2030–2060 under the RCP 8.5 and Melting 1.5 scenarios.

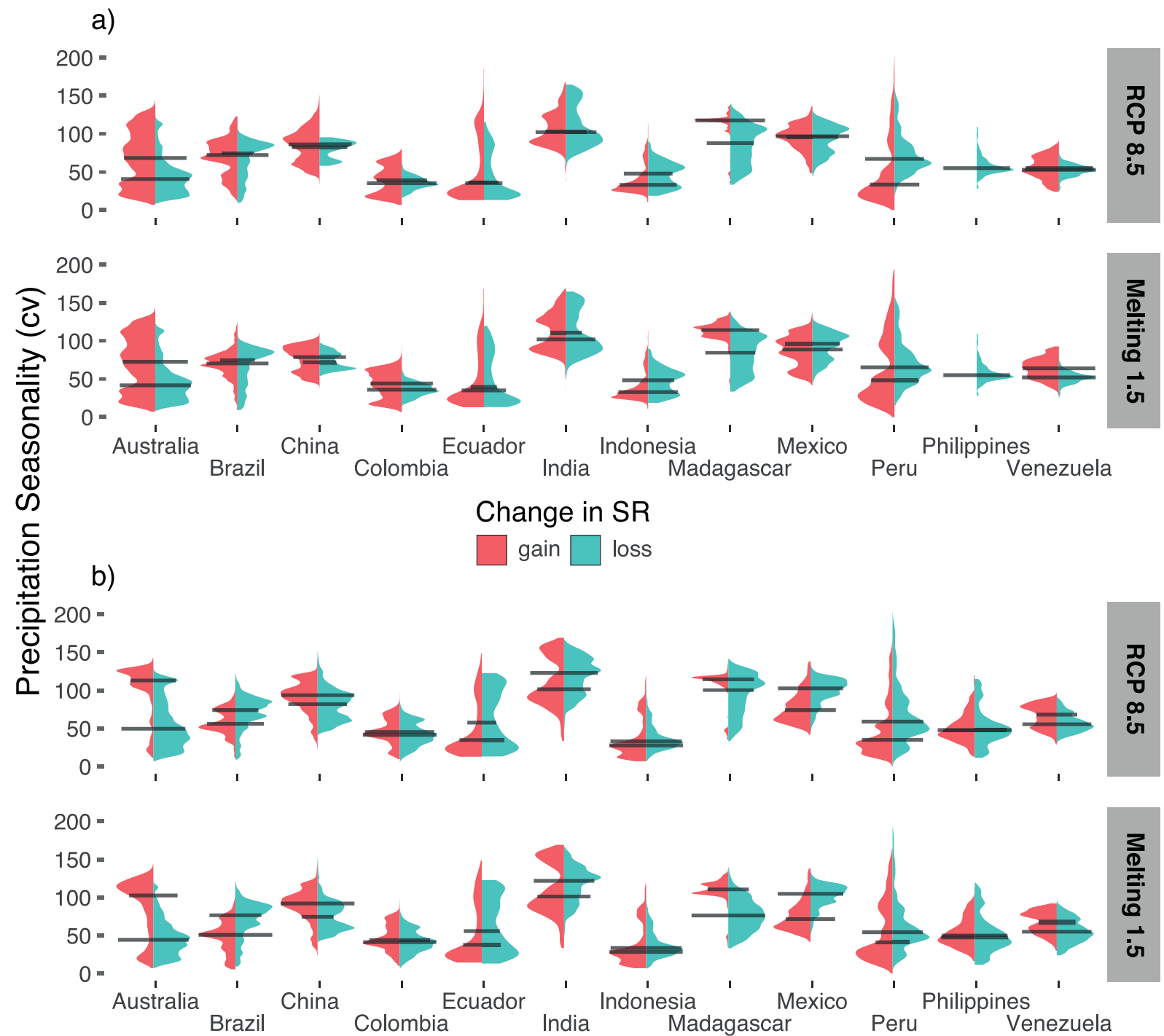

**Supplementary figure S6**

**Present-day precipitation seasonality for areas projected with gain and loss in species richness**

**across twelve megadiverse countries.** a–b, estimates are based on species distribution models (SDMs) of (a) vascular plants and (b) tetrapods. Areas of gain and loss represent pixels with ΔSR above the third quartile and below the first quartile, respectively. The bioclimatic profile for areas is show for 2030–2060 under the RCP 8.5 and Melting 1.5 scenarios.

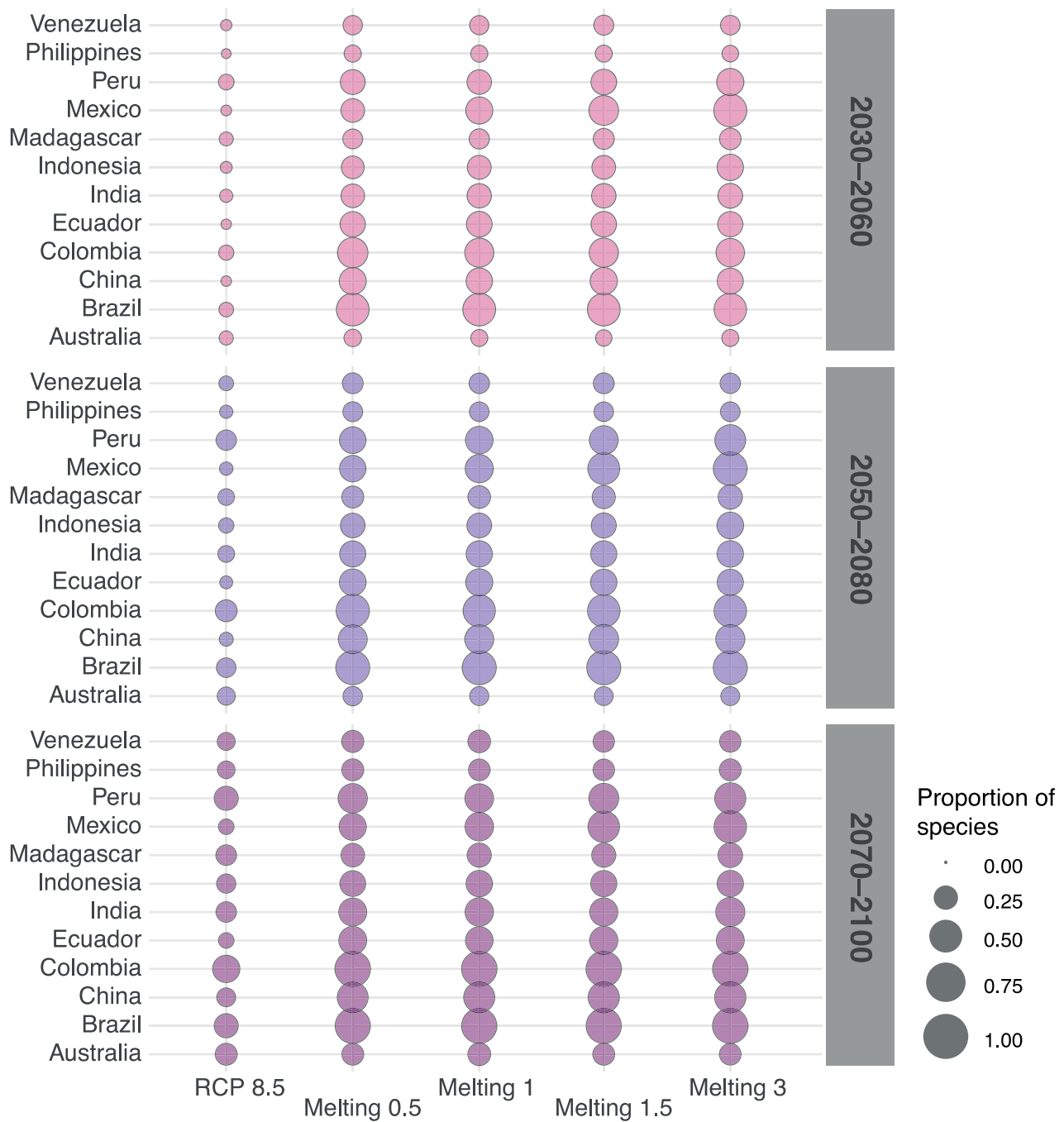

**Supplementary figure S7**

**Proportion of species of vascular plants and tetrapods with complete range loss across twelve megadiverse countries.** The proportion of species (%) with complete range loss was estimated relative to total number of species modelled for each country. Range size estimation was based on species distribution models (SDMs) and were standardized relative to the present-day species range size.

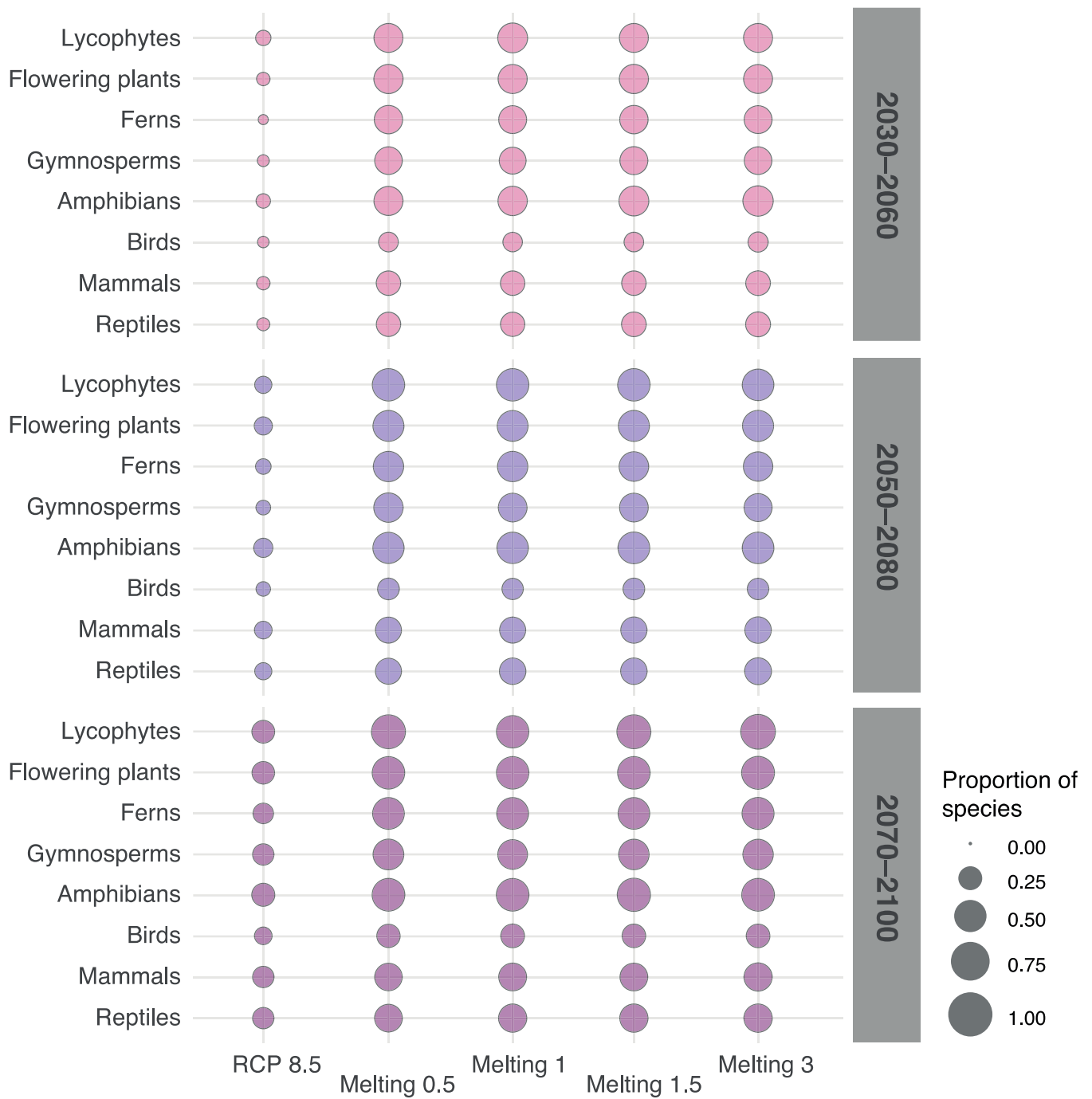

**Supplementary figure S8**

**Proportion of species of vascular plants and tetrapods with complete range loss across eight taxonomic groups.** The proportion of species (%) with complete range loss was estimated relative to total number of species modelled for each country. Range size estimation was based on species distribution models (SDMs) and were standardized relative to the present-day species range size.

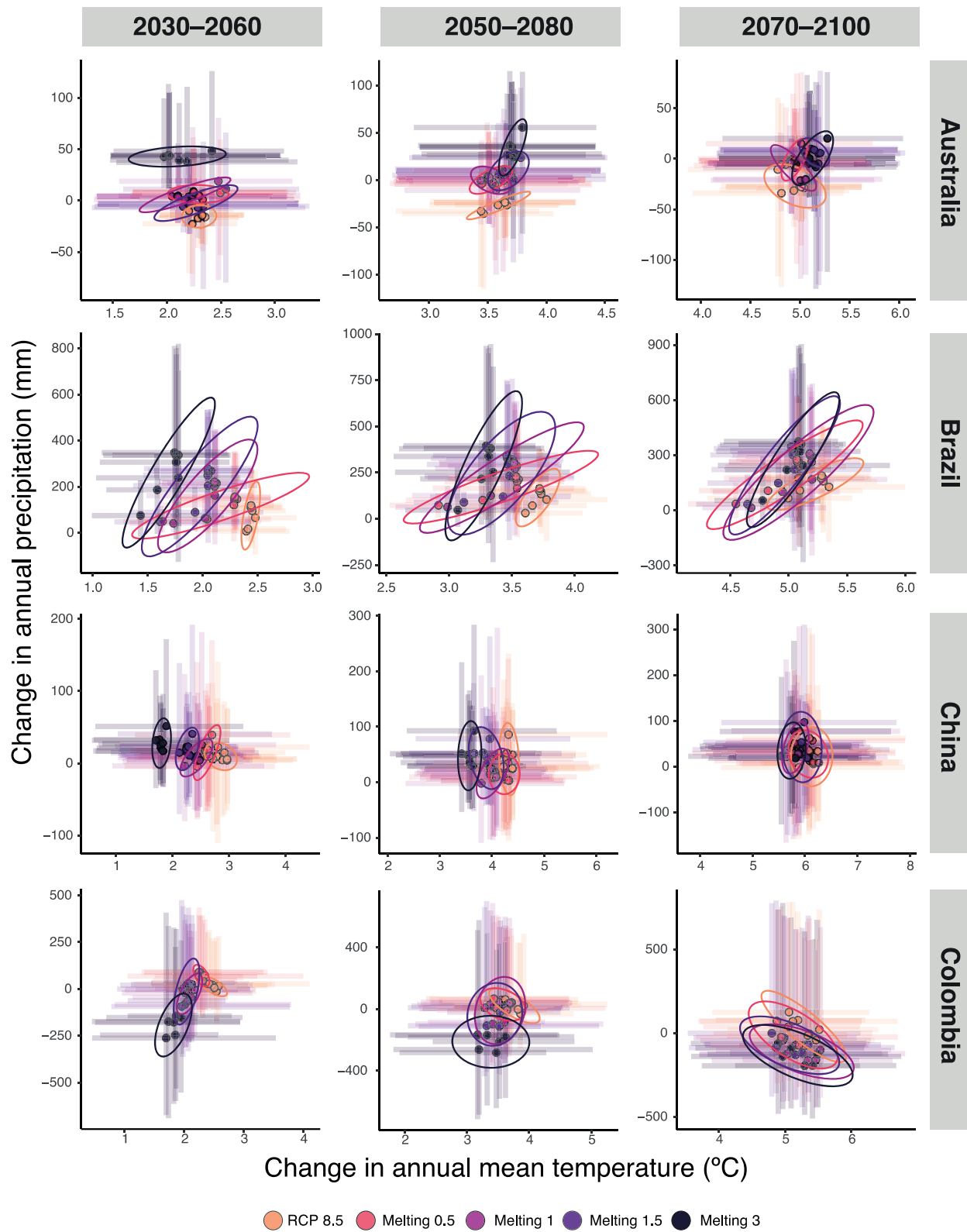

**Supplementary figure S9**

**Temporal changes in mean annual temperature and precipitation projected across megadiverse countries.** Temporal change was estimated as the difference between future climate conditions and

present-day climate conditions across four of the twelve countries. Data are groups by country (rows) and temporal horizon (columns). The different taxonomic groups are depicted by circles and cross lines, which depict the median and the corresponding first and third quartiles of species' values within groups, respectively. Colors correspond to the five future climate models. As a visual guide we provide ellipses around data points for every climate model.

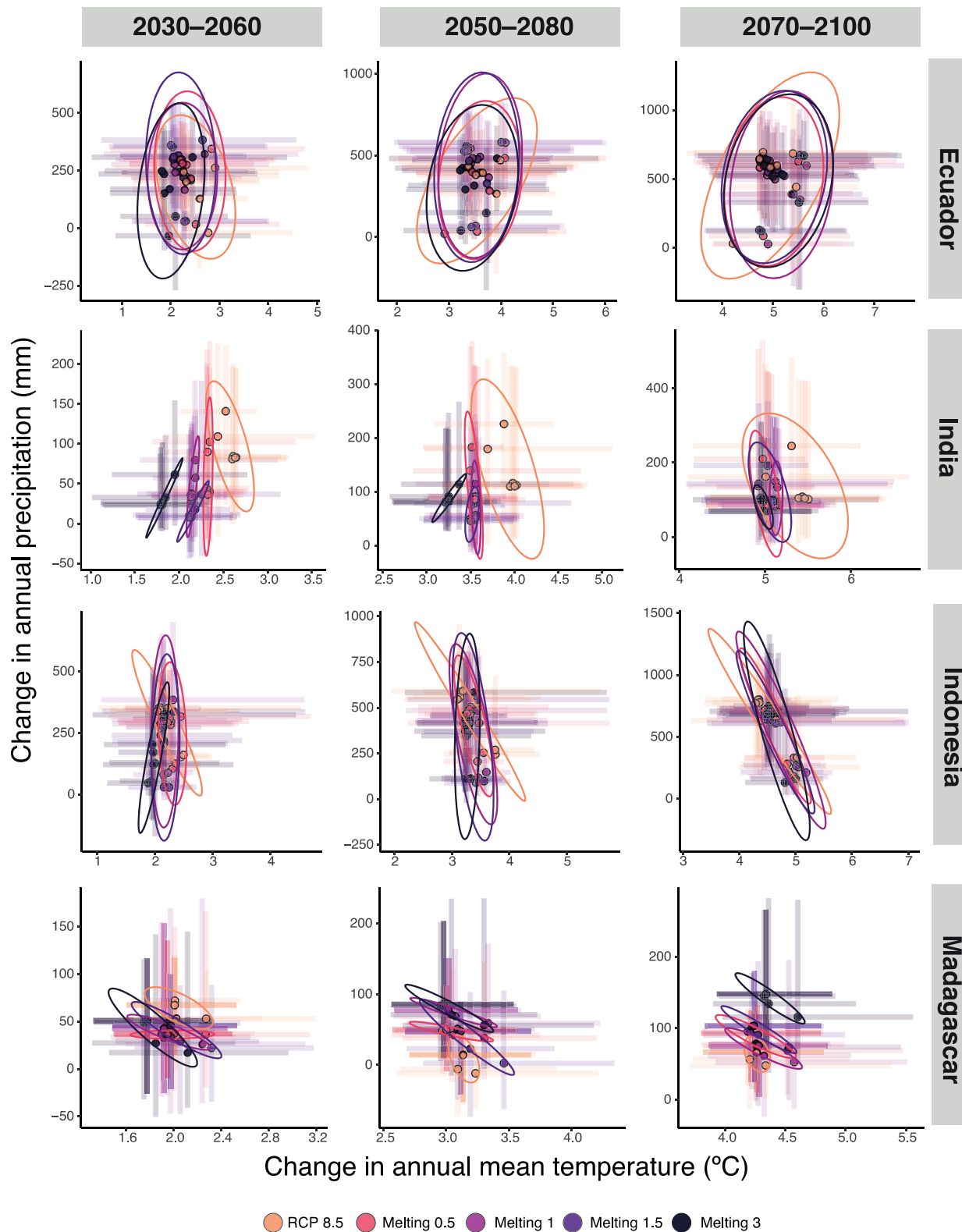

**Supplementary figure S10**

**Temporal changes in mean annual temperature and precipitation projected across megadiverse countries.** Temporal change was estimated as the difference between future climate conditions and

present-day climate conditions across four of the twelve countries. Data are groups by country (rows) and temporal horizon (columns). The different taxonomic groups are depicted by circles and cross lines, which depict the median and the corresponding first and third quartiles of species' values within groups, respectively. Colors correspond to the five future climate models. As a visual guide we provide ellipses around data points for every climate model.

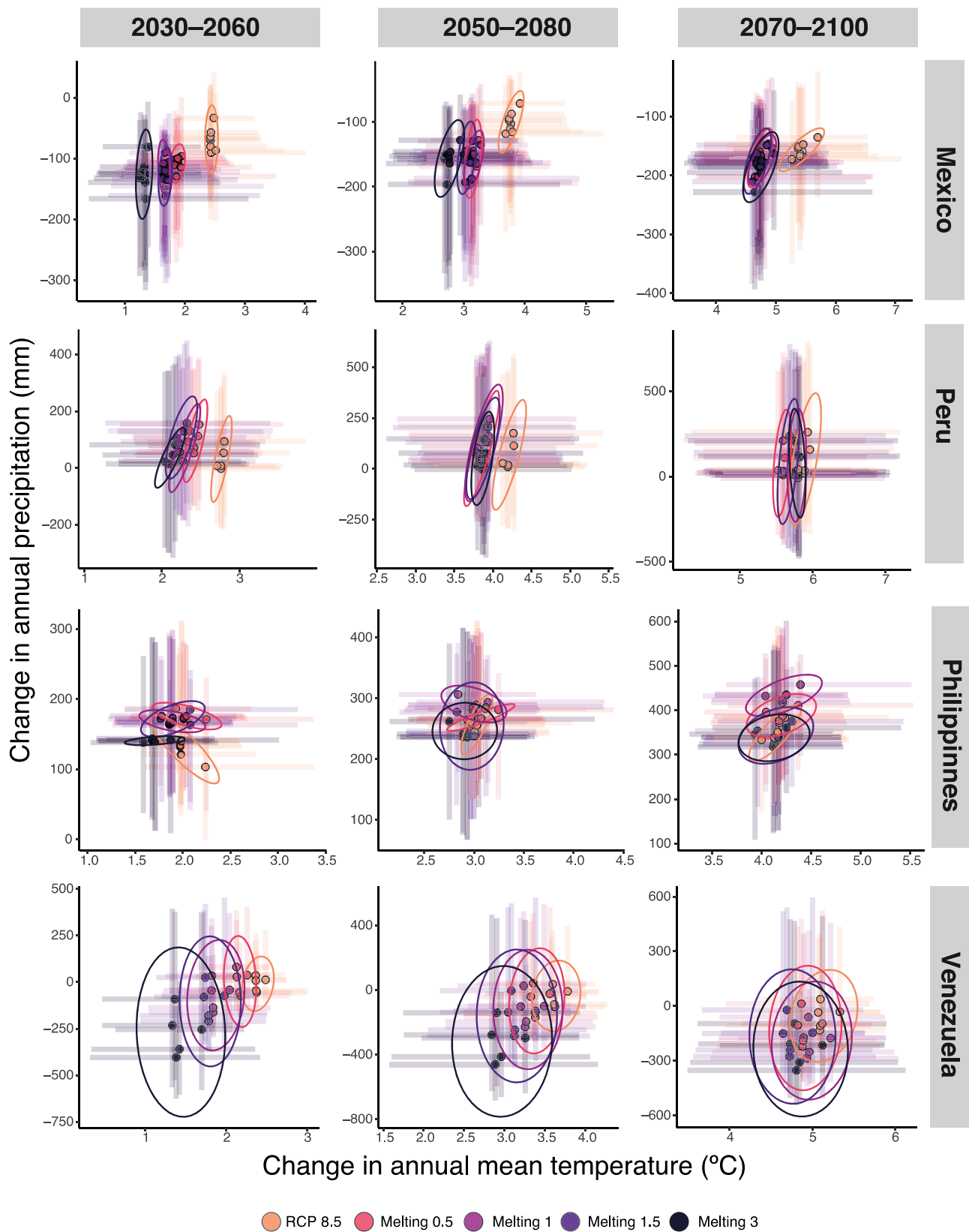

**Supplementary figure S11**

**Temporal changes in mean annual temperature and precipitation projected across megadiverse countries.** Temporal change was estimated as the difference between future climate conditions and

present-day climate conditions across four of the twelve countries. Data are groups by country (rows) and temporal horizon (columns). The different taxonomic groups are depicted by circles and cross lines, which depict the median and the corresponding first and third quartiles of species' values within groups, respectively. Colors correspond to the five future climate models. As a visual guide we provide ellipses around data points for every climate model.

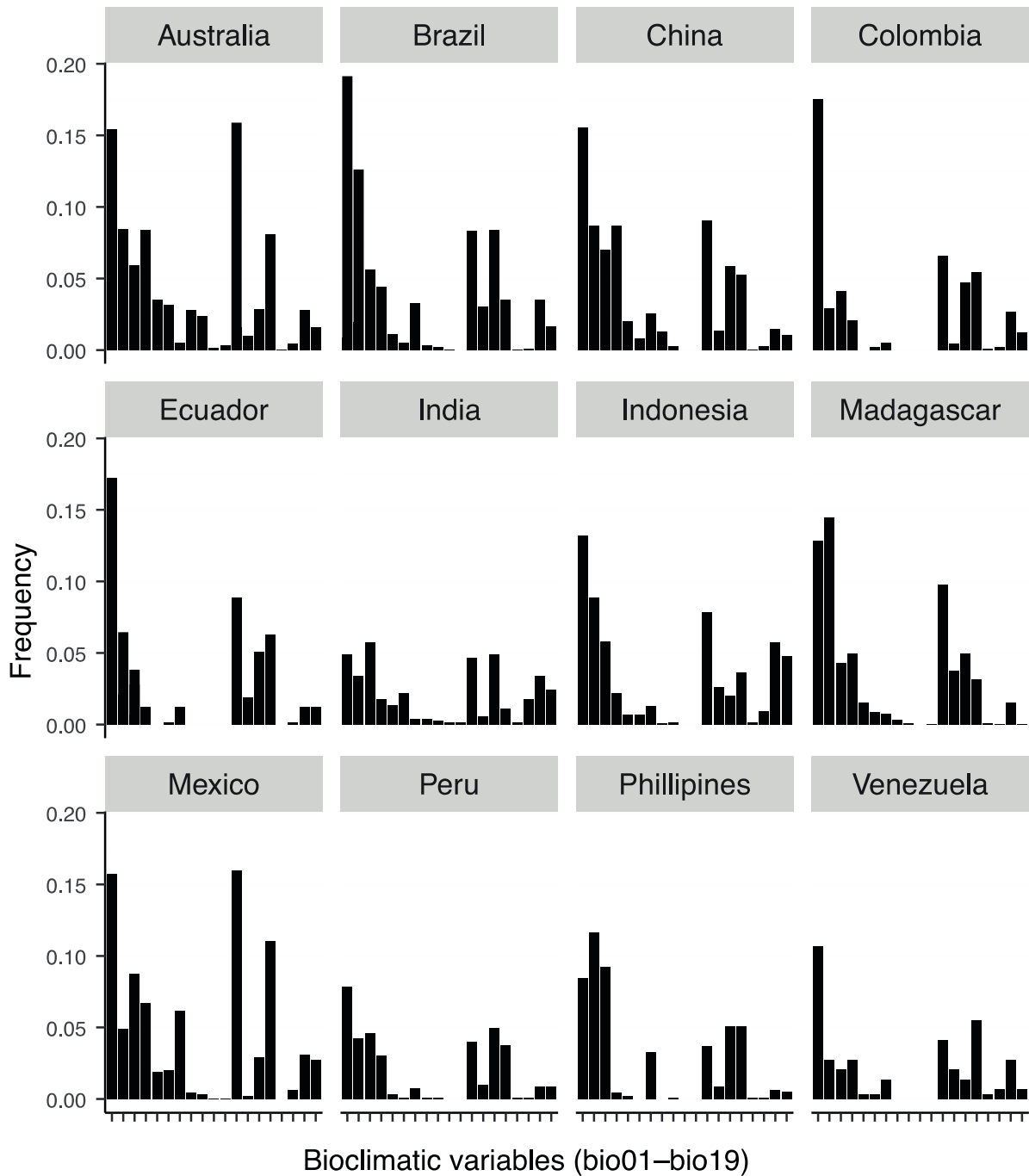

**Supplementary figure S12**

**Bioclimatic variables with the highest contribution to species distribution models.** Frequency of appearance of variables with the highest model contribution across the 21,146 species distribution models grouped by country. Estimates are based on the two variables with the highest contribution for every species model.

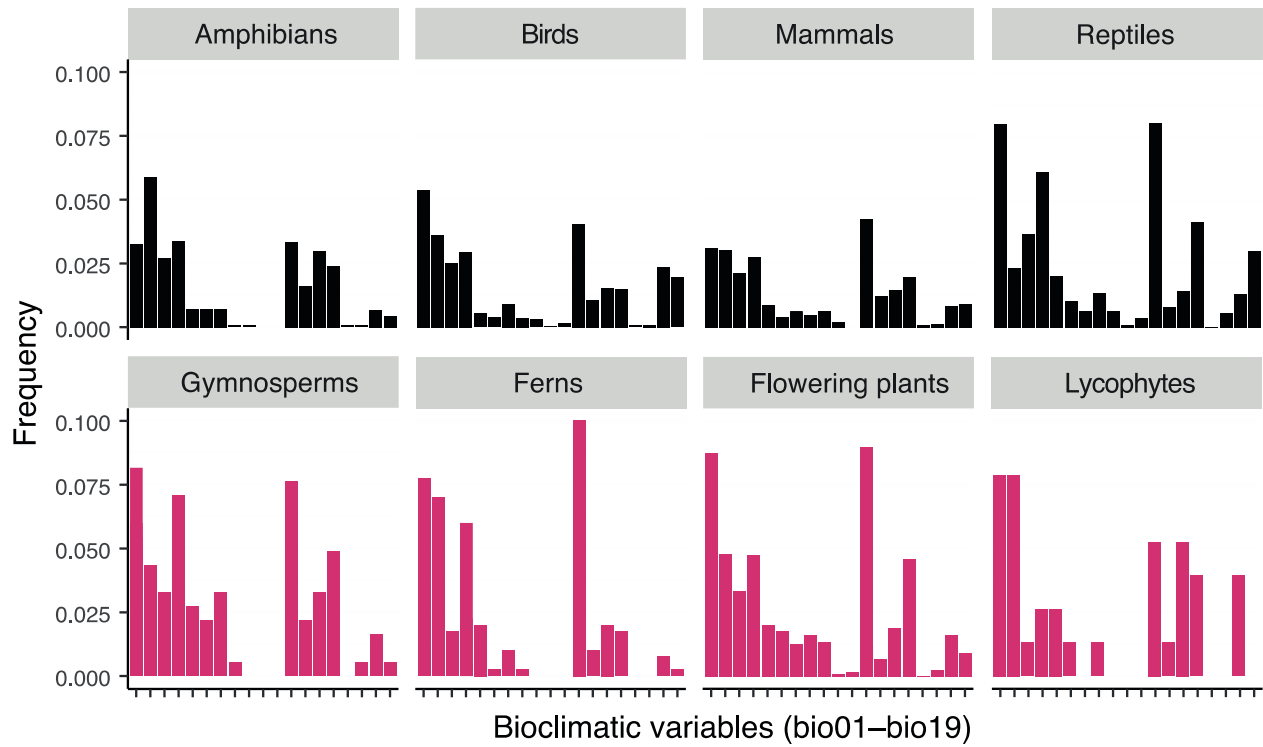

**Supplementary figure S13**

**Bioclimatic variables with the highest contribution to species distribution models.** Frequency of appearance of variables with the highest model contribution across the 21,146 species distribution models grouped by taxonomic group. Estimates are based on the two variables with the highest contribution for every species model.

### Data processing

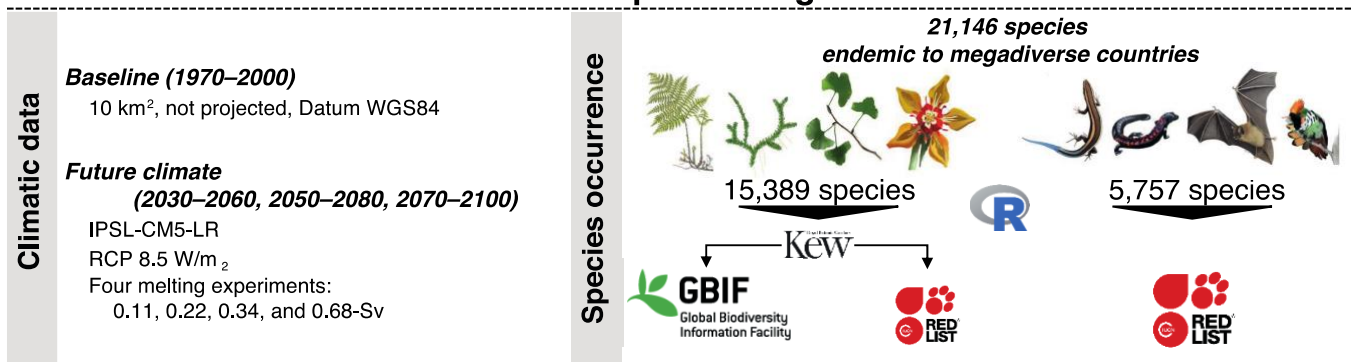

### Species Distribution Modelling

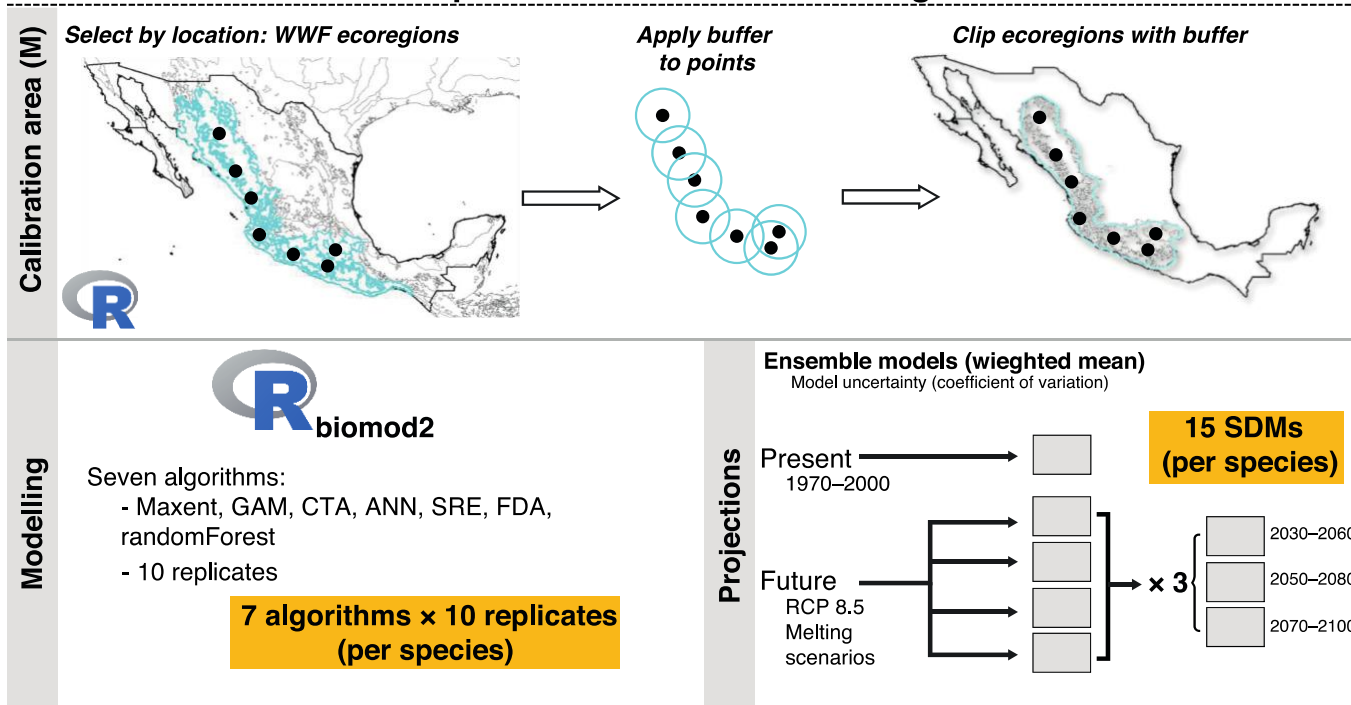

### Spatial analyses

#### Presence/absence matrices (PAM)

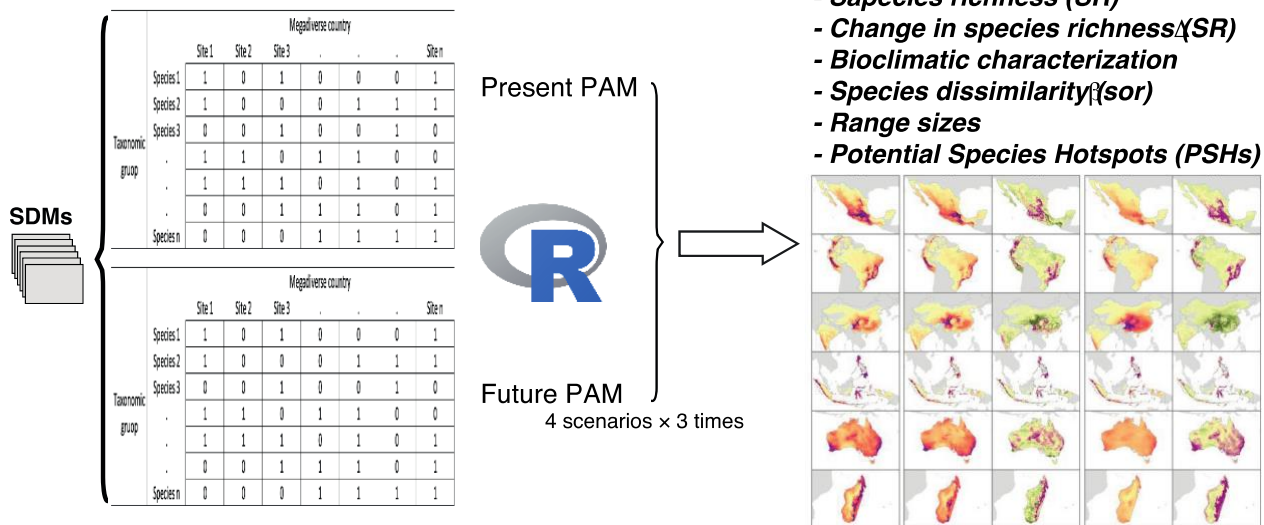

Supplementary figure S14

**Graphical summary of the methodological approach used to model and summarize species distributions.** The diagram depicts the main features of the climatic data and species occurrence records we used. We then briefly show how we selected the calibration areas (M) for every species, which is a critical step during species distribution modelling, describe the algorithms used for modelling, and show the underlying logic to projecting the estimated species' climatic profiles across geography and time. Finally, the diagram depicts the construction of presence/absence matrices from the species distribution models and how these matrices are the basis of all the analyses we performed.

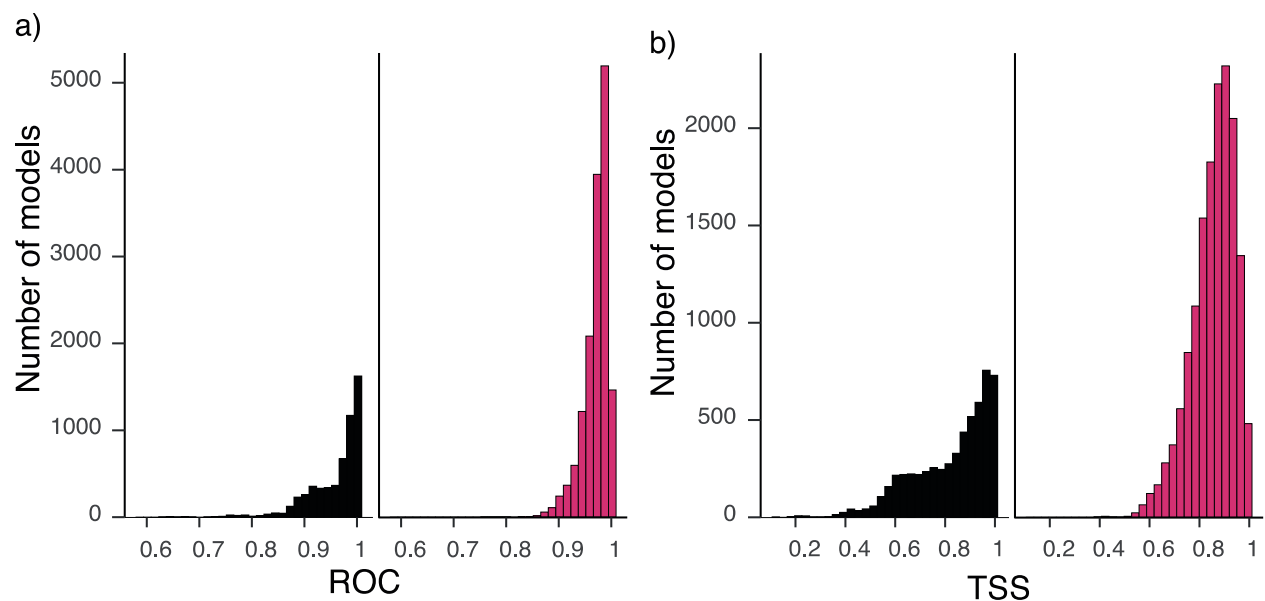

**Supplementary figure S15**

**Validation statistics for the ensemble species distribution models for tetrapods and vascular plants.** a-b, histograms depicting the distribution of (a) ROC and (b) statistics for the 21,146 ensemble models for tetrapods (left panels) and vascular plants (right panels).
